# Precision Repair of Zone-Specific Meniscal Injuries Using a Tunable Extracellular Matrix-Based Hydrogel System

**DOI:** 10.1101/2024.09.12.612723

**Authors:** Se-Hwan Lee, Zizhao Li, Ellen Y. Zhang, Dong Hwa Kim, Ziqi Huang, Sang Jin Lee, Hyun-Wook Kang, Jason A. Burdick, Robert L. Mauck, Su Chin Heo

## Abstract

Meniscus injuries present significant therapeutic challenges due to their limited self-healing capacity and diverse biological and mechanical properties across meniscal tissue. Conventional repair strategies neglect to replicate the complex zonal characteristics within the meniscus, resulting in suboptimal outcomes. In this study, we introduce an innovative, age- and stiffness-tunable meniscus decellularized extracellular matrix (DEM)-based hydrogel system designed for precision repair of heterogeneous, zonal-dependent meniscus injuries. By synthesizing age-dependent DEM hydrogels, we identified distinct cellular responses: fetal bovine meniscus-derived DEM promoted chondrogenic differentiation, while adult meniscus-derived DEM supported fibrochondrogenic phenotypes. The incorporation of methacrylate hyaluronic acid (MeHA) further refined the mechanical properties and injectability of the DEM-based hydrogels. The combination of age-dependent DEM with MeHA allowed for precise stiffness tuning, influencing cell differentiation and closely mimicking native tissue environments. *In vivo* tests confirmed the biocompatibility of hydrogels and their integration with native meniscus tissues. Furthermore, advanced 3D bioprinting techniques enabled the fabrication of hybrid hydrogels with biomaterial and mechanical gradients, effectively emulating the zonal properties of meniscus tissue and enhancing cell integration. This study represents a significant advancement in meniscus tissue engineering, providing a promising platform for customized regenerative therapies across a range of heterogeneous fibrous connective tissues.

## Introduction

The region-specific composition and architecture of fibrous connective tissues, such as tendons, ligaments, and the meniscus, present significant challenges for effective repair and regeneration. These tissues are characterized by complex biochemical compositions and biomechanical properties that are not uniform, but rather vary across different regions.[1] For instance, a gradient of collagen types and the spatial distribution of proteoglycans endow different areas within the same single tissue with distinct mechanical characteristics.[2] This complexity is further compounded by the presence of diverse cell types, each finely tuned to their specific microenvironment.[3] This zonal heterogeneity is essential for the proper function of these tissues, but also complicates the development of effective treatment strategies. As a consequence, a traditional ‘one-size-fits-all’ approach to implant design is likely to be ineffective, emphasizing the need for targeted therapies that can be tuned to reflect the intricate variations in these native tissues.

The meniscus is an important example of a heterogeneous tissue, as the diversity in biochemical and biomechanical properties is crucial for knee joint biomechanics, facilitating load-bearing and uniform force distribution in the knee. Meniscal tears, often caused by congenital defects, degenerative changes, or sports-related factors, impact roughly 70 individuals per 100,000 annually[4] and often require surgical intervention due to the limited self-repair capacity of the tissue.[5] The meniscus is distinctly zonal: the outer (red-red) region is vascularized, harboring fibroblast-like cells and type-I collagen which imparts stiffness and durability to the tissue (**Figure 1a**). This outer region is particularly stiff and resistant to tensile forces due to its robust collagen structure.^[3a]^ In contrast, the inner (white-white) region is avascular, populated by chondrocyte-like cells, and rich in type-II collagen and proteoglycans, with a more randomly organized collagen fiber network (**Figure 1a**).[6] This inner zone is more compressible and prone to deformation compared to the outer and middle zones.[7] The intermediate (red-white) zone serves as a transitional area, blending characteristics of both the outer and inner regions.[8] Moreover, these zonal differences necessitate repair strategies that are tuned to each zone: the outer zone has better healing potential due to its vascularity, while the inner zone poses a greater challenge for repair due to its avascularity.[9]

**Figure 1.**
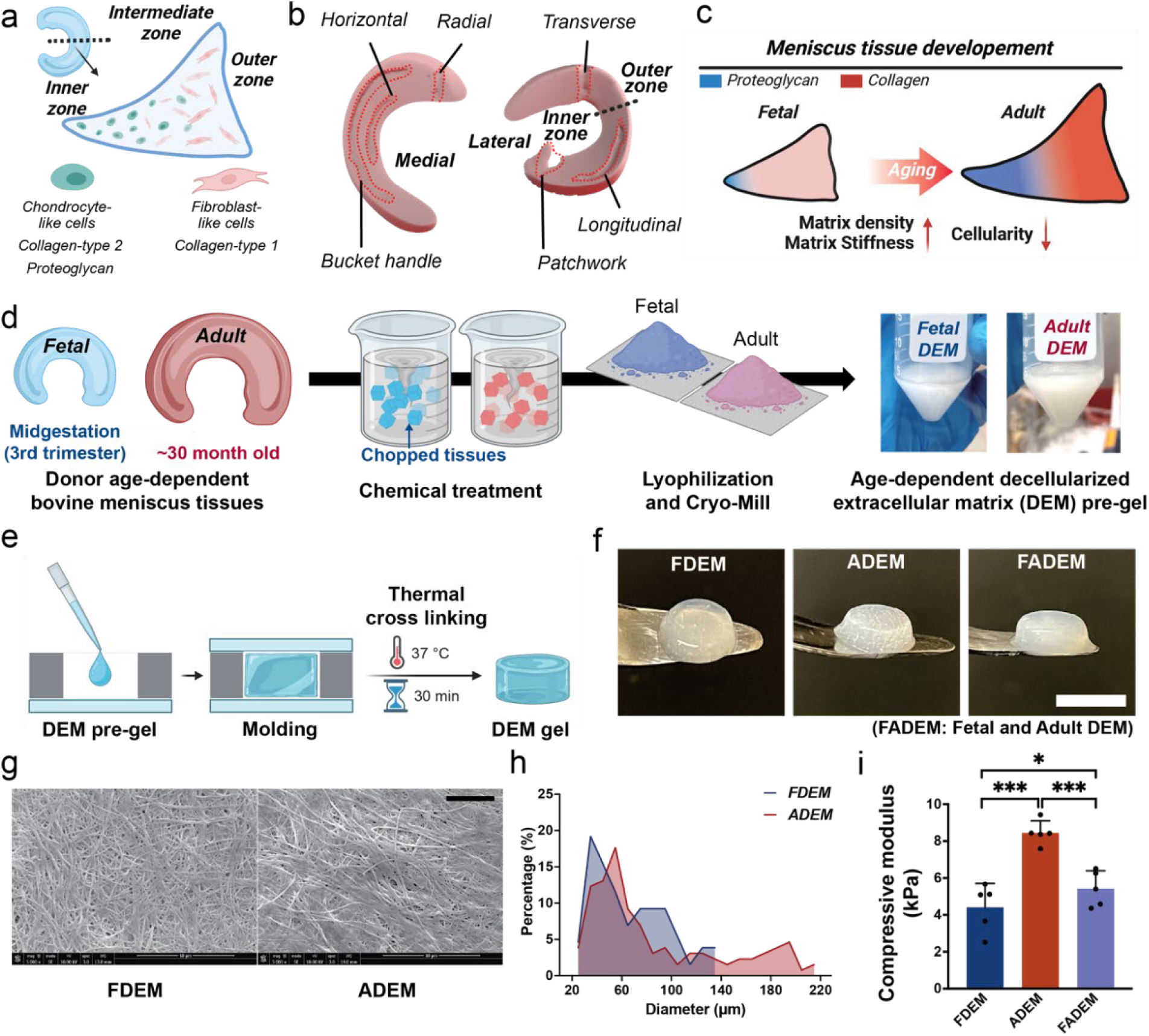
Decellularization and Characterization of Age-Dependent Meniscus Decellularized Extracellular Matrix (DEM) Hydrogels. Schematics showing a) heterogenous cell types and biochemical compositions across different meniscus zones, b) different types of meniscus tears that vary in direction and location, and c) biochemical alterations in meniscus tissue during development. Schematics of d) the SDS-based decellularization process to generate age-dependent DEM hydrogel precursors (fetal: FDEM, adult: ADEM) and e) DEM hydrogel fabrication process using molding and thermal crosslinking. Characterization of DEM hydrogels, including f) images of fabricated DEM hydrogels (scale bar: 5 mm), g) field emission SEM images of the fibrous microstructure of DEM hydrogels (scale bar: 5 µm), h) fiber size distribution in FDEM and ADEM hydrogels, and i) compressive moduli of FDEM, ADEM, and their combination (FADEM) hydrogels (n=5 per group; *p<0.05, ***p<0.001).

Meniscus tears also exhibit significant heterogeneity and variety (e.g., location, direction), with common types including longitudinal, radial, and horizontal tears (**Figure 1b**).[9] These tears, each with their distinct shape, orientation, and affected zones, underscore the necessity for customized treatment strategies.[10] These strategies must be sensitive to the unique attributes of each tear type to optimize healing and restore knee functionality. Commonly employed techniques such as suturing and arthroscopic partial meniscectomy (APM) offer contrasting approaches;[11] suturing aims to preserve and mend the tissue, while APM involves excising the damaged segment.[12] Despite their widespread use, both methods exhibit notable shortcomings when confronting complex meniscus tears: suturing is often inadequate for complex tear patterns (e.g., radial tears),[13] and APM results in increased local contact pressures that contribute to more rapid joint degeneration or osteoarthritis.[14] This highlights a significant gap in clinical practice, pointing to the urgent need for more sophisticated, tailored therapeutic modalities that ensure effective healing and joint preservation.

Recently, decellularized extracellular matrix (DEM) systems have emerged as promising substrates for meniscus repair, offering a conducive microenvironment for cellular proliferation and tissue regeneration.[15] Yet, the challenge remains of effectively emulating the zonal-dependent properties and intrinsic heterogeneity of the meniscus using such systems. This challenge calls for innovative strategies that leverage DEM systems to mimic features of the distinct biochemical and biomechanical profiles of various zones in the meniscus. Therefore, the goal of this study was to engineer a tunable meniscus DEM-based hydrogel system that caters to the zonal specificity of meniscus tissue. By customizing the biochemical and mechanical properties of these hydrogels, we sought to mimic the native tissue characteristics of each meniscal zone. Moreover, the meniscus undergoes significant changes in protein composition and mechanical properties during development (**Figure 1c**).[16] This is evident when comparing fetal (from the midgestation, second and third trimester) and adult (from skeletally mature, 20-30 months) bovine menisci, which demonstrate an age-related increase in collagen and proteoglycan content from the inner to the outer zone.[17] Notably, donor age significantly influences tissue size and mechanical stiffness.[18]

On this basis, here, we leveraged age-specific meniscus DEM — FDEM from fetal and ADEM from adult donors — to create biomimetic hydrogel systems. Our comprehensive biological characterizations confirmed that the ECM components and mechanical properties of the hydrogel, which are donor-age-dependent, indeed modulate meniscus cell behaviors and phenotypes. Building upon these insights, we furthered tuned the properties of DEM-based injectable hydrogel systems, by incorporating stiffness-tunable methacrylate hyaluronic acid (MeHA),[19] allowing precise control over hydrogel stiffness. MeHA also enhanced injectability and printability, expanding the potential routes to apply these hydrogels within surgical treatments. These advanced hydrogels, designed to mimic aspects of native ECM, offer promising zone-specific treatment strategies for fibrous connective tissue injuries, advancing tissue engineering and optimizing patient outcomes.

## Results

### Development and Characterization of Age-Dependent Meniscus DEM Hydrogels

Considering the influence of meniscal development on the biochemical and mechanical properties of the tissues,[20] which in turn may subsequently affect cellular phenotypes, we first established an age-dependent DEM system. This system utilizes bovine menisci of varying age (fetal: FDEM, adult: ADEM) to explore the regulatory effects of the age-dependent meniscus ECM on cell behavior. Specifically, FDEM and ADEM from bovine tissue was generated through an SDS-based decellularization process (**Figure 1d**). Post-decellularization, effective removal of the majority of cells was confirmed via DAPI and H&E staining (**Supplementary Figure S1a-d**). Cylindrical DEM hydrogels (Diameter: 5 mm, Thickness: 2 mm) were produced using a molding method coupled with thermal crosslinking (**Figure 1e-f**). Additionally, a composite group, FADEM, was created by combining FDEM and ADEM at a 1:1 ratio.

Scanning electron microscopy (SEM) analysis verified the preservation of fibrous structures in both FDEM and ADEM hydrogels (**Figure 1g**). Notably, ADEM hydrogels exhibited a broader fiber diameter distribution compared to FDEM hydrogels, likely reflecting developmental changes in ECM composition (**Figure 1h**).[21] The compressive modulus of the ADEM hydrogel (8.44 ± 0.65 kPa) was double that of the FDEM hydrogel (4.41 ± 1.29 kPa). Blending these two DEM types (FADEM) prior to gel formation (**Figure 1i**) resulted in a compressive modulus (5.42 ± 0.96 kPa) that was intermediate to the FDEM and ADEM hydrogels alone. These distinctive features, including differences in fibrous structure and enhanced modulus, suggest that the age-related characteristics of native tissue are retained post-decellularization.

### Impact of Age-Dependent Meniscus DEM on Cell Response

Based on our expectation of age-dependent differences in the ECM of menisci that form these hydrogel systems, we hypothesized that these differences would translate into distinct cellular responses. Thus, we next explored how variations in the DEM composition, reflective of different developmental stages, affected cellular behavior, particularly with mesenchymal stem cells (MSCs) and meniscus fibrochondrocytes (MFCs). To do this, both cell types were first cultured on age-dependent DEM hydrogels (**Figure. 2a**). MSCs cultured on ADEM hydrogels exhibited a large, more elongated cell morphology, while those on the FDEM hydrogel were notably smaller and more rounded (**Figures 2b-d**, **Supplementary Figure S2a**). MSCs on DEM hydrogels that combined features of both fetal and adult stages (FADEM) displayed intermediate morphologies. Similar trends were observed in MFCs under analogous conditions (**Supplementary Figure S3a-e**), suggesting that the age-specific mechanical properties and biochemical cues of the DEM significantly influence cell morphology and phenotype across cell types.

**Figure 2.**
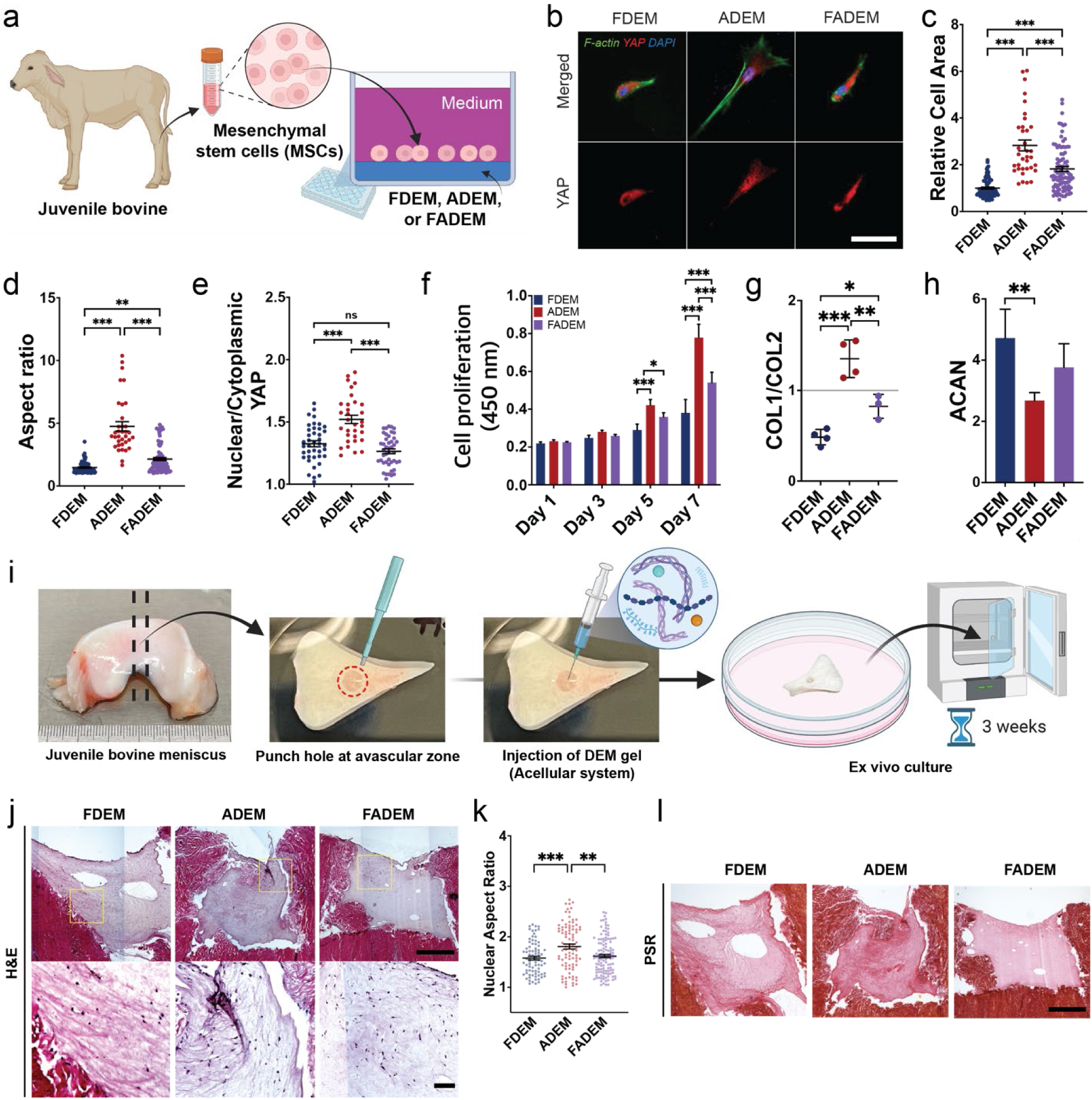
Influence of Meniscus Age on Cellular Interactions with DEM hydrogels. a) Schematic of *in vitro* evaluation of cell response to DEM hydrogels from different ages (FDEM, ADEM, FADEM). Characterization of MSC interactions with DEM hydrogels at day 3, including b) representative images (Green: F-actin, Red: YAP, Blue: DAPI; Scale bar: 50 µm), c) relative cell area (n=35-87), d) aspect ratio (n=35-87), and e) YAP nuclear localization (n=32-44). Characterization of MSCs with longer term culture on DEM hydrogels (for 7 days), including f) cell proliferation (n=5) and g-h) gene expression (n=3-4, COL1/COL2 and ACAN) (*p<0.05, **p<0.01, ***p<0.001). i) Schematic of implantation and culture for 3 weeks of acellular DEM hydrogels within meniscus explants and outcomes, including j) representative H&E images [Scale bars: 750 µm (top) and 100 µm (bottom)], k) quantified nuclear aspect ratio of recruited cells (n=67-118; **p<0.01, ***p<0.001), and l) representative picrosirius red stained images (Scale bar: 750 µm).

To further assess cellular mechano-response, Yes-associated protein (YAP) localization was quantified. In both FDEM and FADEM hydrogels, YAP was predominantly localized to the cytoplasm, whereas in the ADEM hydrogel, YAP concentration was notably higher in the nucleus (**Figure 2b and e**). This differential localization indicates distinct cellular responses to mechanical cues in their environment: nuclear YAP presence in the ADEM hydrogel suggests activation of pathways favoring fibrochondrocyte differentiation, essential for the load-bearing function of the outer meniscus[22], while cytoplasmic YAP in the FDEM and FADEM hydrogels may reflect a softer ECM that promotes chondrogenic differentiation[23].

Indeed, MSCs and MFCs both demonstrated significantly higher proliferation rates when cultured on the ADEM hydrogel (**Figure 2f and Supplementary Figure S3f**). Notably, FDEM hydrogel cultures exhibited a pronounced upregulation of chondrogenic markers, including Aggrecan (ACAN), SRY-Box Transcription Factor 9 (SOX9), and Transforming Growth Factor (TGF), while ADEM hydrogel cultures showed increased expression of fibrogenic markers including Collagen type I alpha 2 chain (Col1a2) in MSCs (**Figure 2g and h**, **Supplementary Figure S2b**). The ratio of the Col1a2 to Collagen type II (Col2) was elevated in the ADEM compared to FDEM hydrogel, suggesting a phenotype closer to fibrocartilage-like tissue in MSCs cultured on the ADEM hydrogel and chondrogenic tissue in MSCs cultured on the FDEM hydrogel.[24] The FADEM hydrogel presented intermediate characteristics in terms of cell proliferation and gene expression for both cell types (**Figure 2f-h** and **Supplementary Figure S2b**). Similarly, in MFCs, chondrogenic expression (e.g., ACAN, SOX9, and TGF) was upregulated for cells on FDEM hydrogels, while fibrogenic expression, including Col1a2 and Connective Tissue Growth Factor (CTGF), was more prominent for cells on ADEM hydrogels (**Supplementary Fig. S3g**). These findings demonstrate that the FDEM microenvironment fosters a chondrogenic phenotype and augments expression pertinent to tissue remodeling, whereas ADEM facilitates cell proliferation and a fibrogenic phenotype.

### Long-Term Assessment of Age-Specific Meniscus DEM Hydrogels in Ex Vivo Models

Given that long-term stability and endogenous cell recruitment are crucial for successful tissue regeneration with hydrogels[25], we next assessed the performance of the age-dependent meniscus DEM hydrogels within a physiologically relevant environment. Specifically, we introduced acellular hydrogels into an ex-vivo meniscus defect model and incubated for three weeks (**Figure. 2i**). Hematoxylin and eosin (H&E) staining verified the preservation of FDEM, ADEM, and FADEM hydrogels throughout the incubation period, along with substantial cell infiltration from the surrounding native tissue (**Figure. 2j**). Consistent with our *in vitro* observations, cells migrating into the ADEM hydrogels exhibited more cell nuclei compared to those in other groups (**Figure. 2j, k**). All hydrogel types preserved their collagen composition, with ADEM hydrogels demonstrating more intense collagen staining (**Figure 2l**). The persistence of collagen suggests that these hydrogels maintain their structural integrity over time, providing a stable scaffold conducive to tissue regeneration.

### Proteomic Analysis of Age-Dependent DEM Hydrogels Reveals Distinct Biochemical Composition

Given the observed differences in cellular responses to age-dependent DEM hydrogel compositions and the known impact of tissue development on the biochemical composition of the meniscus[16], we performed a comprehensive proteomic analysis to investigate these differences. Samples were obtained from four distinct fetal and adult meniscus donors, and subsequent principal component analysis (PCA) and heat map generation of detected proteins revealed minimal variability between individual donors, resulting in the formation of four distinct clusters (**Figure 3a and b**, **Supplementary Figure S5a and b**).

**Figure 3.**
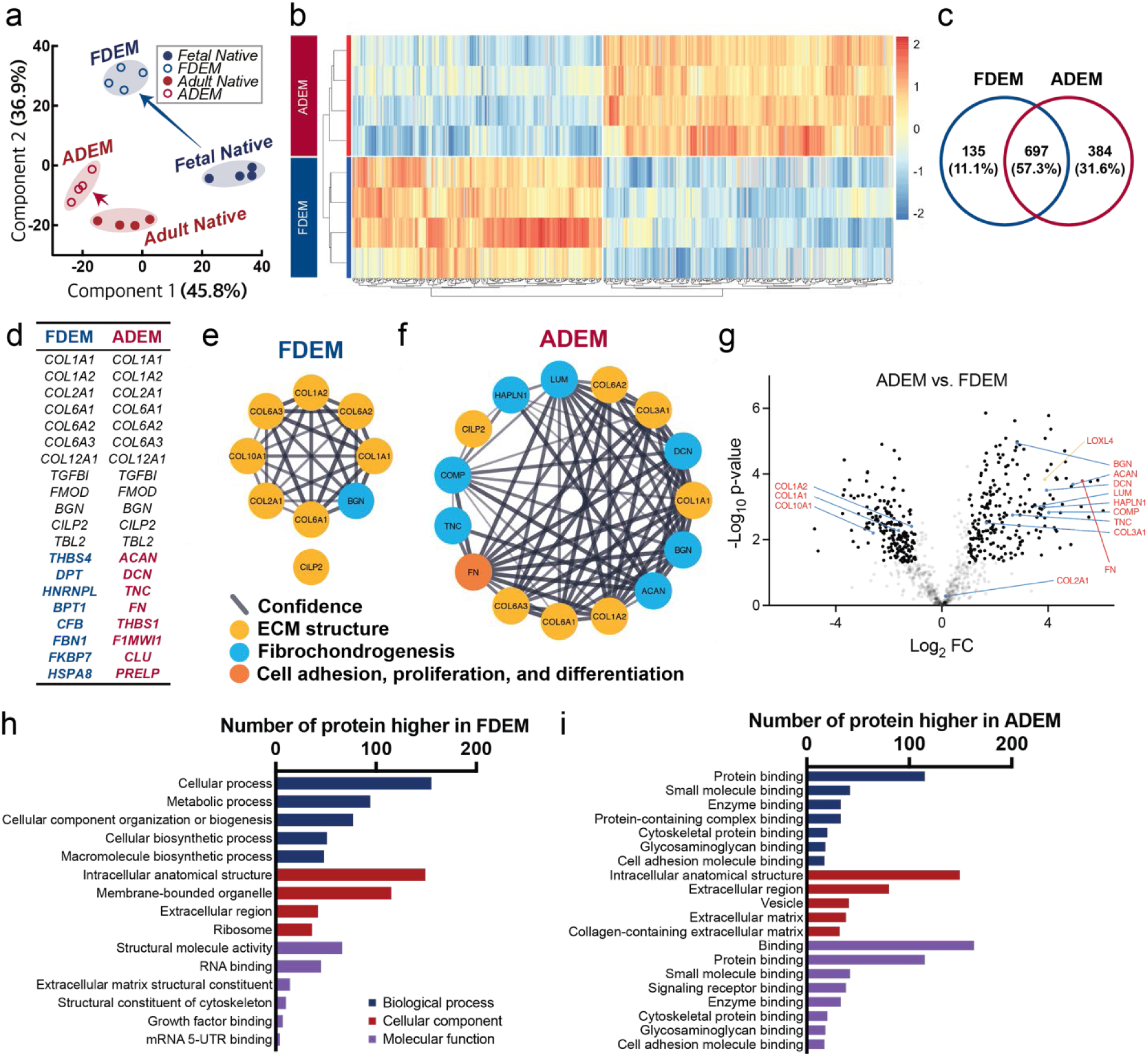
Proteomic Analysis of Age-Dependent Meniscus DEM. Proteomic analysis of FDEM and ADEM derived from four different donors per group, including a) principal component analysis (PCA) showing the distribution and variance of the protein composition, highlighting the distinct proteomic landscapes of FDEM and ADEM, b) heatmap and clusters of detected proteins, with clusters indicating the proteomic similarities and differences between the groups, c) Venn diagram depicting the overlap and unique proteins preserved in FDEM and ADEM, d) top 20 most prevalent proteins in FDEM or ADEM, Protein-protein interaction networks related to ECM structure, fibrochondrogenesis, cell adhesion, proliferation, and differentiation in (e) FDEM and (f) ADEM, with line thickness representing the confidence level of data support. g) Volcano plot of ADEM (vs. FDEM) visualizing the differentially expressed proteins between ADEM and FDEM, with statistical significance and fold-change metrics. Gene ontology analysis for biological process, cellular component, and molecular function in (h) FDEM and (i) ADEM showing the top 8 categories.

The analysis identified 1,081 proteins in adult-derived DEM (ADEM) compared to 832 proteins in fetal-derived DEM (FDEM), indicating a more diverse and abundant array of proteins in ADEM (**Figure 3c**). Post-decellularization analysis confirmed the retention of several proteins integral to the native meniscus in both the FDEM and ADEM. These include Collagen Type-1 (COL1), Collagen Type-2 (COL2), Collagen Type-3 (COL3), Collagen Type-6 (COL6), Collagen Type-12 (COL12), Biglycan (Bgn), and Cartilage Intermediate Layer Protein 2 (CILP2) (**Figure 3d and Supplementary Figure S5c**). Notably, COL6 was abundant in both FDEM and ADEM. In addition to these structural proteins, our proteomic analysis revealed a significant presence of proteins associated with wound healing, regulatory functions, and ECM assembly. Proteins such as Transforming growth factor-beta-induced protein ig-h3 (TGFBI), Transducin beta like 2 (TBL2), and fibromodulin (FMOD) were abundant in both FDEM and ADEM **(Figure 3d)**.[26] A more intriguing aspect revealed by proteomic analysis was the prevalence of fibrochondrogenic proteins in ADEM compared to FDEM. Fibrochondrogenic proteins such as Aggrecan (ACAN), Decorin (DCN), and Tenascin C (TNC) were more prevalent in ADEM than FDEM. Remarkably, Fibronectin (FN), a protein pivotal to cell adhesion and proliferation, was exclusively identified in ADEM group, where it exhibited significant interactions with other structural proteins, namely collagens **(Figure 3d-g)**. The retention of these major proteins post-decellularization suggests that essential components of native tissue are largely preserved after the process **(Supplementary Figure S5)**.

In addition, a volcano plot further highlighted differentially abundant proteins in ADEM compared to FDEM, showing that proteins associated with fibrochondrogenesis (e.g., ACAN, DNC, and TCN) and cell adhesion (e.g., FN) were expressed at higher levels in ADEM than in FDEM (**Figure 3e**). This differential expression likely contributes to the enhanced mechanical properties of ADEM hydrogels. Furthermore, Lysyl oxidase-like 4 (LOXL4), a protein implicated in collagen crosslinking[27] and fibrosis[28], was significantly upregulated in ADEM (**Figure 3e**). These expression patterns align with those observed in native meniscus tissues (**Supplementary Figure S5d, e**).

The abundance of cell adhesion-related proteins in ADEM may create a more favorable environment for cellular attachment to the structural proteins compared to FDEM. This environment may encourage cells to adopt larger and elongated shapes, supporting cell proliferation and fibrochondrogenesis. The protein networks identified post-decellularization also form interconnected network structures similar to native tissue, potentially providing cells with a microenvironment that closely resembles the native ECM. Additional gene ontology (GO) analysis revealed that FDEM was enriched with proteins associated with cellular and metabolic processes, while ADEM exhibited a higher prevalence of proteins associated with protein or ECM binding (**Figure 3h, i and Supplementary Figure S4a, b**). Notably, GO analysis identified that ADEM is enriched in proteins related to cytoskeletal protein binding and cell adhesion molecule binding (**Figure 3i**). The GO categories in DEM hydrogels closely align with those in native tissues, indicating that the decellularization process effectively preserves key protein compositions (**Supplementary Figure S5f, g**).

In summary, the proteomic analysis of FDEM and ADEM reveals critical biochemical differences that likely drive the distinct cellular responses to these age-dependent systems. The retention of key structural and functional proteins, particularly in ADEM, underscores its potential to emulate the fibrochondrogenic native meniscus environment effectively.

### Decoupling DEM Hydrogel Composition and Stiffness on Cellular Response

Given that FDEM and ADEM hydrogels had distinct stiffnesses and biochemical compositions, we next sought to determine which of these factors drove the cellular response. To address this, we doubled the concentration in FDEM (2×FDEM) to achieve a compressive modulus comparable to that of ADEM hydrogels (**Figure 4a**), standardizing the stiffness between the two systems. Interestingly, despite this adjustment, MSCs on 2×FDEM hydrogels still had a lower proliferation rate than those on ADEM (**Figure 4b**). Moreover, cells on the 2×FDEM were smaller and more circular compared to their counterparts on ADEM hydrogels (**Figure 4c-f**). This was further corroborated by significantly reduced YAP nuclear localization in MSCs on the 2×FDEM compared to ADEM hydrogels (**Figure 4g**). These findings suggest that stiffness alone does not fully account for the varied cellular responses seen in age-dependent DEM hydrogels, underscoring the importance of biochemical cues.

**Figure 4.**
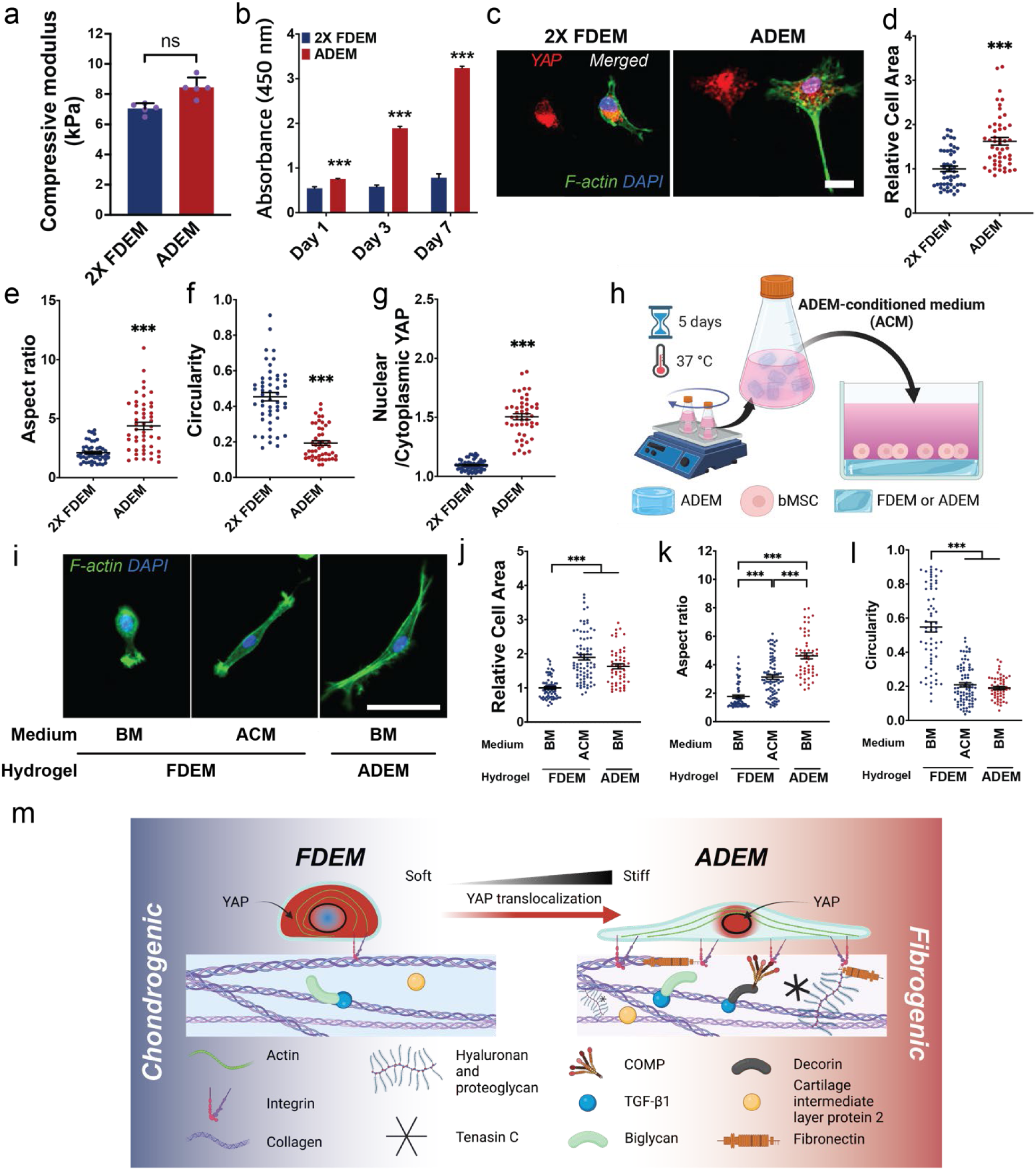
Biochemical and Mechanical Effects of DEM Hydrogels. a) Compressive modulus of 2×FDEM and ADEM hydrogels (n=5; ns: not significant). b) MSC proliferation on 2×FDEM and ADEM hydrogels at days 1, 3, and 7 (n=5; ***: p<0.001 vs. 2X FDEM). MSC morphology analysis after 3 days, including c) representative F-actin and YAP staining (Scale bar: 20 µm), d) relative cell area (n=49), e) aspect ratio, f) circularity, and g) YAP nuclear localization (n=46-57; ***p<0.001 vs. 2×FDEM). h) Schematic of study where releasate from ADEM is added to MSC cultures atop FDEM or ADEM hydrogels. Cell morphology under varied media and hydrogel conditions: i) Representative F-actin images (Scale bar: 50 um). MSC morphology analysis, including j) relative cell area, k) aspect ratio, and l) circularity (n=52-82; ***p<0.001). m) Summary schematic showing the different biochemical and biomechanical microenvironments of the FDEM and ADEM hydrogels that drive chondrogenic and fibrogenic differentiation, respectively.

To further investigate the impact of biochemical composition, ADEM hydrogels were dissolved in basal growth media to prepare ADEM-conditioned media (ACM). MSCs were seeded on either FDEM or ADEM hydrogels and cultured with ACM or BM (as a control group, **Figure 4h**). Interestingly, ACM culture prompted cells on FDEM hydrogels to enlarge and elongate, resembling the morphology of cells on ADEM hydrogels under BM conditions (**Figure 4i-l**). Similar results were observed with MFCs cultured on tissue culture plates (TCP) (**Supplementary Figure S6a-d**) and ACM did not compromise cell viability (**Supplementary Figure S6b**). Additionally, super-resolution H2B STORM imaging demonstrated that ACM culture reduced nanoscale chromatin condensation in MFC nuclei (**Supplementary Figure S6e and f**), suggesting enhanced transcriptional activation in MFCs. Furthermore, DTAF and PSR staining confirmed that MFCs cultured in ACM produced more extracellular matrix and collagen than those in the control group (**Supplementary Figure S6g-i**). These findings highlight the pivotal role of biochemical composition over mechanical stiffness in dictating cellular response to DEM hydrogels. The increased cell proliferation and morphological changes observed with ACM imply that specific biochemical factors inherent to ADEM are key drivers of these responses. The unique protein profiles revealed by proteomic analysis, particularly those associated with cell adhesion and ECM composition, may create a favorable microenvironment for cell adhesion, growth, and differentiation.

Taken together, this differential enhancement is crucial for meniscus tissue engineering, where regeneration of specific tissue types and functions is required. Notably, the fibrochondrogenic biochemical cues from ADEM alongside its stiffer biomechanical environment, significantly enhance cell attachment, proliferation, and YAP nuclear localization. These factors collectively foster an environment favorable to fibrochondrogenic differentiation and the formation of fibrous cartilage. Conversely, the biochemical signals and softer biomechanical properties of FDEM favor chondrogenesis (**Figure 4m**). These insights underscore the importance of biochemical signals in the design of tissue engineering scaffolds for meniscus repair and regeneration, highlighting the need for a tunable approach in scaffold design that integrates both mechanical and biochemical considerations to optimize regenerative outcomes.

### Biochemically and Mechanically Tunable DEM-based Hydrogels for Targeted Zone-specific Meniscus Repair

Given the complexity of the meniscus, injectable materials tailored to these specific meniscus zones are needed. Building from the zonal-specific cues provided by dECM of different ages, we next developed a stiffness-tunable DEM-based injectable hydrogel system. For this, we combined our DEM with methacrylated hyaluronic acid (MeHA), whose mechanics can be adjusted by controlling the degree of crosslinking (**Figure 5a**).[29] To first generate a ‘Soft’ MeHA and ‘Stiff’ MeHA component, the methacrylation level was set at 22% and 100%, respectively (**Supplementary Figure S7a, b**). ‘Soft’ MeHA had a compressive modulus of 3.1 (± 0.4) kPa while ‘Stiff’ MeHA was 16.2 (± 2.3) kPa (**Supplementary Figure S7c**). Next, the age-dependent DEM and MeHA were combined in predetermined ratios (**Figure 5a**). Fluorescence imaging (z-stack; z-axis distance: 200 μm) using FITC-labeled MeHA confirmed a homogeneous mixture of the two components (**Supplementary Figure S7e and Video 1**). The resulting hydrogels exhibited a wide range of stiffness, from 4.4 kPa (FDEM) to 27.3 kPa (ADEM/Stiff MeHA) (**Fig. 6b and Supplementary Figure S7d**). By mixing ‘Soft’ and ‘Stiff’ at a 1:1 ratio, hydrogels with an intermediate stiffness were generated. Picrosirius red and Alcian blue staining confirmed that collagen content was preserved in the DEM and that proteoglycans were replenished from the added MeHA (**Supplementary Figure S7f, g**). These results demonstrated that, by modifying the methacrylation degree of HA, we could create DEM-based hydrogels with a broad range of stiffness values, to create tailored microenvironments for meniscus repair, potentially supporting the regeneration of specific tissue types and functions.

**Figure 5.**
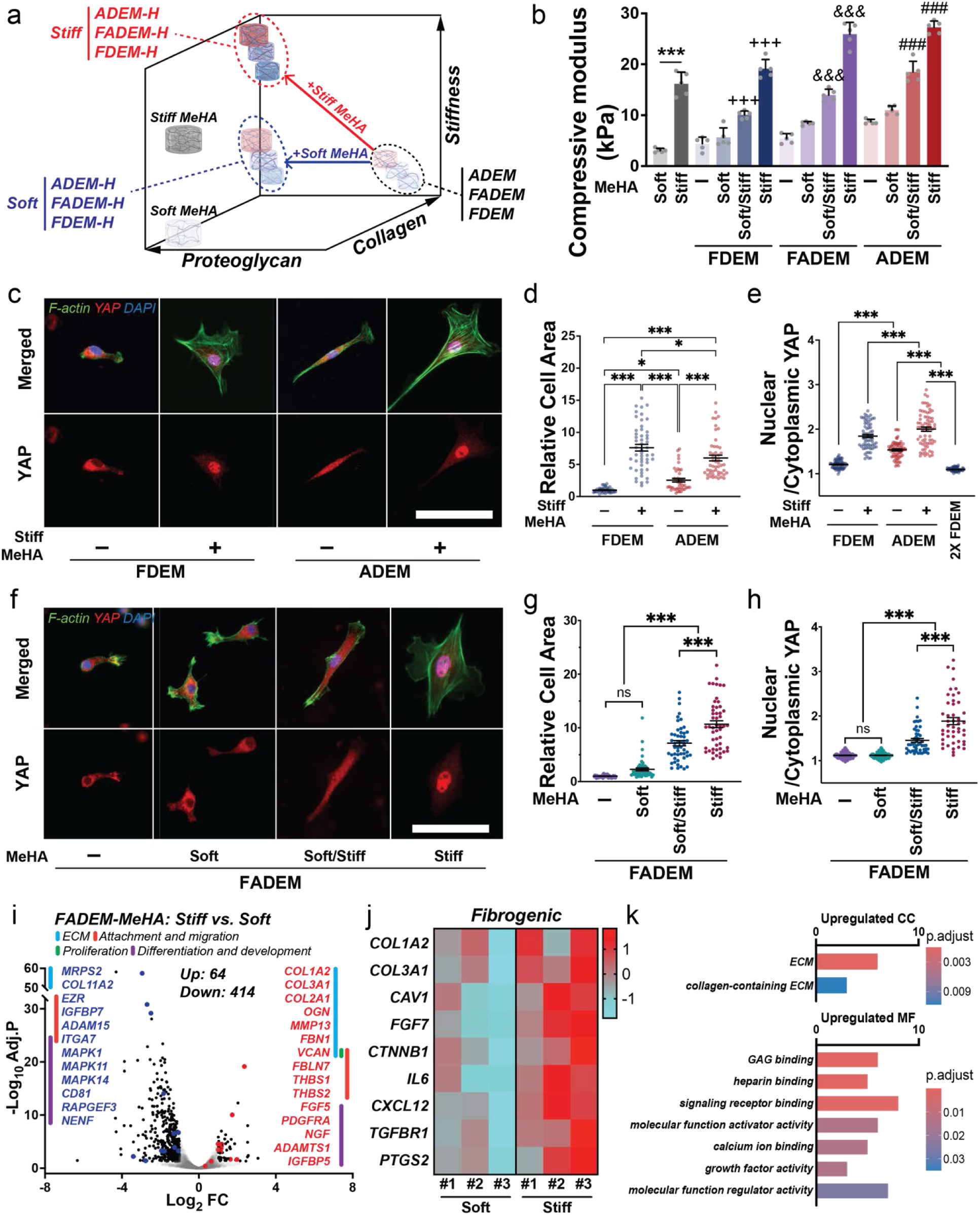
Stiffness-tunable DEM-Based Hydrogels Direct MSC Phenotype. a) Illustration of strategies to develop tunable DEM-based hydrogel systems for zonal-specific meniscus tissue repair by including methacrylated hyaluronic acid (MeHA) of varied stiffness. b) Stiffness of mechanically modulated DEM-based hydrogels (n=5; ***: p<0.001, +++: p<0.001 vs. FDEM, &&&: p<0.001 vs. FADEM, ###: p<0.001 vs. ADEM, p values for all comparison cases in **Supplementary Table 2**). MSC response to age-dependent DEM-based hydrogels at day 3, including c) representative images on age-dependent DEM-based hydrogel systems with or without the addition of ‘stiff’ MeHA (Green: F-actin, Red: YAP, and Blue: DAPI), d) relative cell area (n=42-50; *p<0.05, ***p<0.001), e) YAP nuclear localization (n=67-72; ***p<0.001). MSC response to stiffness tunable FADEM-based hydrogels after 3 days, including f) representative images on FADEM hydrogels with or without ‘soft’, ‘soft/stiff’, or ‘stiff’ MeHA (Green: F-actin, Red: YAP, and Blue: DAPI), g) relative cell area (n=50; *p<0.05, ***p<0.001), and h) YAP nuclear localization (n=50; ***p<0.001). Transcriptional analysis to compare MSCs on FADEM-Stiff MeHA (Treatment) vs. FADEM-Soft MeHA (Control) via RNA sequencing (n=3/group), including i) Volcano plot, j) Z-score heat maps of fibrogenic gene expression, k) GO analysis of gene differentially expressed genes (CC: cellular components and MF: molecular functions).

**Figure 6.**
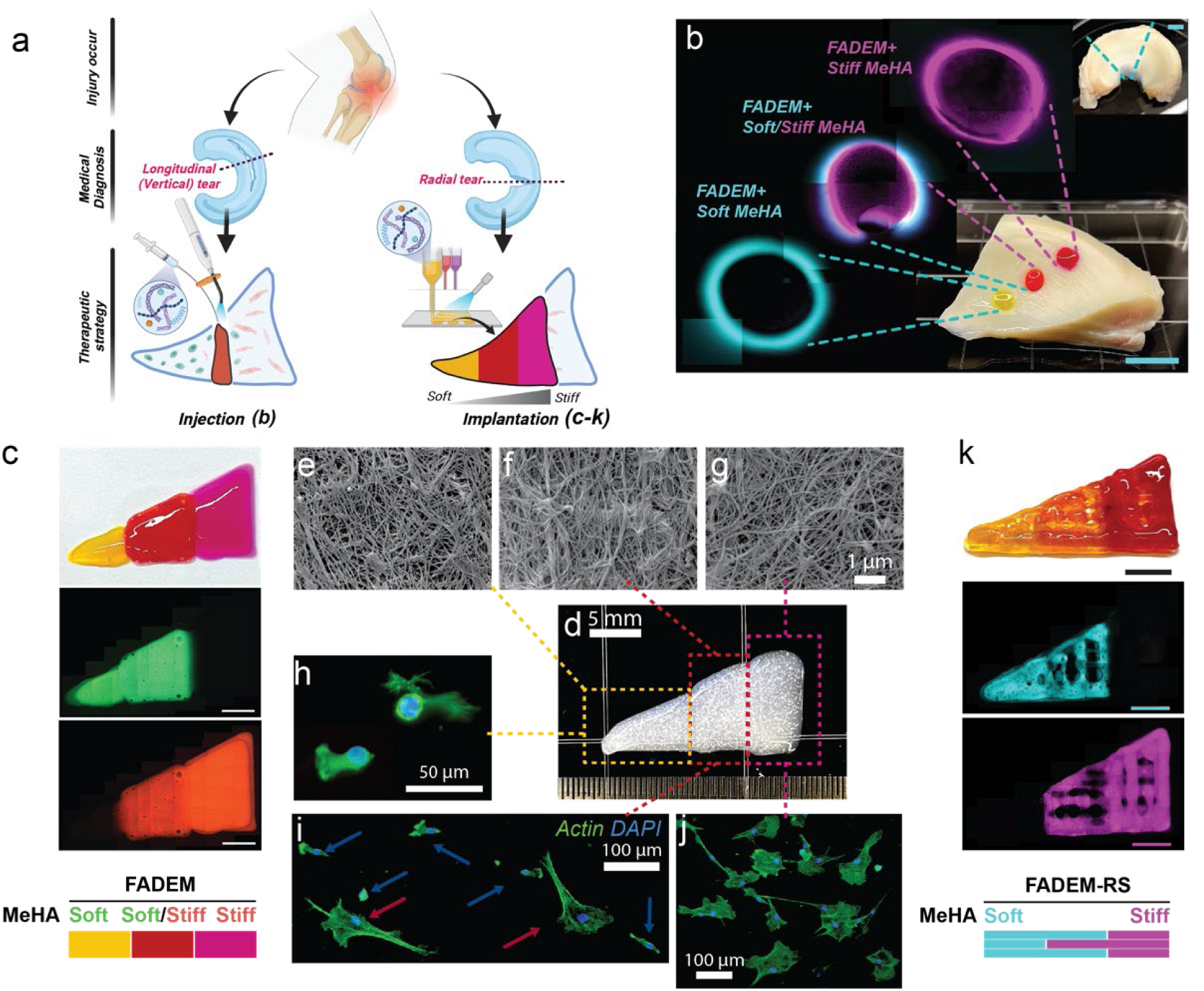
Targeted Meniscus Repair Using Stiffness-Tunable DEM-Based Hydrogels. a) Illustration of the application of the system. *Injection method*: b) injected stiffness-modulated FADEM-based hydrogel in defects in different meniscus zones (scale bar: 5 mm). *Implantation method using 3D printed hybrid constructs*: c) images of a bioprinted stiffness-modulated FADEM-based hydrogel (yellow: printed ‘soft’ FADEM hydrogel for the inner meniscus zone, red: ‘soft/stiff’ FADEM hydrogel for the middle meniscus zone, purple: ‘stiff’ FADEM hydrogel for the outer meniscus zone, scale bar: 5 mm). (e-g) Representative FE-SEM images of fibrous structure of printed FADEM-based hydrogels. (h-j) F-actin staining of MSCs cultured on the zone-dependent printed FADEM-based hydrogels. (k) 3D printed lattices of zone-specific FADEM-based hydrogels with the addition of ruthenium/sodium persulfate (FADEM-RS, cyan: ‘soft’ FADEM, magenta: ‘stiff’ FADEM, scale bars: 5 mm).

### Cellular Responses in Tunable DEM-based Hydrogel Systems

Next, to determine how the presence of MeHA and the stiffness of tunable DEM-based hydrogels impacted cellular response, MSCs were cultured on the FDEM or ADEM hydrogels with or without the addition of ‘stiff’-MeHA. On these ‘stiff’ DEM-based hydrogel systems, incorporation of ‘stiff’ MeHA into age-dependent hydrogels significantly increased cell area and YAP nuclear localization compared to FDEM, ADEM, or 2×FDEM (which has a similar stiffness to the ADEM) hydrogels (**Figure 5c-e**).

Given that the FADEM hydrogel (a 1:1 mix of FDEM and ADEM) had mechanical and cellular characteristics intermediate between FDEM and ADEM hydrogels and contains both FDEM and ADEM proteins, we further examined cellular responses on FADEM-based stiffness-modulated MeHA hydrogels. When MSCs were cultured on FADEM-based MeHA hydrogels, cells on the ‘Soft’ hydrogels remained small and circular but elongated (**Figure 5f-h**, **Supplementary Figure S7j**) and increased YAP nuclear localization (**Figure 5f, i**) with increased modulus. Notably, ‘Stiff’ FADEM-based hydrogel exhibited heightened COL1A2 expression, while the chondrogenic markers SOX9 and TGF increased in all MeHA-supplemented groups (**Supplementary Figure S7j**). These data indicate that the introduction of MeHA to DEM elevated chondrogenic gene expression and that ‘stiff’ MeHA increased fibrochondrogenic gene expression in MSCs (**Supplementary Figure S7j**).

To further understand changes in transcriptional profiles induced by the stiffness-tuned FADEM-based hydrogels, we performed whole-transcriptome analysis on two experimental conditions: ‘Soft’ and ‘Stiff’ FADEM-based hydrogels. Unsupervised clustering and PCA visualization (**Supplementary Figure S8a**) showed that samples were clustered based on the stiffness of MeHA. After filtering for significant changes, genes were divided into two distinct clusters (**Supplementary Figs. S8b**). Applying thresholds (adjusted *p* < 0.05, −2 > Fold change (FC), and FC > 2) identified 64 up-regulated and 414 down-regulated genes (**Figure 5i and Supplementary Table 2**). Notably, in the ‘stiff’ FADEM-based microenvironment, meniscus ECM production, cell adhesion, cell migration, and cell development-related genes were upregulated (**Figure 5i**). Further, heatmaps of differential gene expression confirmed that representative fibrogenic genes such as COL1A2 and COL3A1 were significantly expressed in the ‘stiff’ FADEM environment with upregulations of chondrogenic genes including COL2A1, ACAN, and TGFβ3 (**Figure 5i** and **Supplementary Figure S8c**). In contrast, genes related to chondrogenesis, including Cartilage oligomeric matrix protein (COMP) were more predominately expressed in ‘soft’ FADEM environment (**Supplementary Figure S8c**). Moreover, the ‘stiff’ FADEM-based system enhances gene expression related to cell adhesion, such as FN1 (**Supplementary Figure 8d**). These findings confirm that the ‘soft’ FADEM hydrogels promote a chondrogenic phenotype, whereas ‘stiff’ FADEM drives cells towards a fibrocartilage phenotype.

GO analysis provided a more in-depth understanding of the biological processes involved: ECM-related cellular component genes were significantly upregulated in the ‘stiff’ FADEM group (**Figure 5k**). Additionally, GO analysis revealed that genes related to the binding of major proteins including, glycosaminoglycan (GAG) and heparin, and cellular activities were significantly upregulated in the ‘stiff’ FADEM hydrogels (**Figure 5k**). Those data indicate that the ‘stiff’ FADEM microenvironment appears to accelerate cellular and ECM protein production and maturation compared to the ‘soft’ FADEM microenvironment. Taken together, these findings demonstrate that by fine-tuning the stiffness of FADEM-based hydrogels, we can create tailored microenvironments that influence MSC behavior and phenotype.

Finally, to further evaluate the biocompatibility and integration of tunable FADEM-based hydrogel *in vivo*, bovine meniscus explants were prepared by filling FADEM-based hydrogels (**Figure S9a**) and implanted subcutaneously into mice (**Figure S9b**). Three weeks post-implantation, the implants were fully integrated with the host tissue of the mouse (**Figure S9c**). No adverse tissue reaction was observed in any of the groups and histological results were consistent across all groups (**Figure S9d-e**). In the ‘soft’ FADEM group, a considerable number of cells infiltrated the hydrogel, while in the ‘stiff’ FADEM group, a small number of cells were observed migrating at the junction of the scaffold and hydrogel in the central region, forming a thin layer. This phenomenon could be attributed to the differences in the stiffness and degradability of the hydrogels, corroborating the cell infiltration and migration observed in animal experiments.

### Application of Stiffness-Tunable DEM-based Hydrogels for Precision Meniscus Repair

The tunable properties of the stiffness-tunable DEM-based hydrogel system offers a promising avenue for the treatment of diverse meniscus tears by virtue of their tailored the biochemical and biomechanical properties. For example, in the case of either small or longitudinal (vertical) tears within a specific region, a single injection could prove effective, where hydrogels with varying degrees of stiffness — ranging from ‘soft’, to ‘soft/stiff’, to ‘stiff’— could be strategically administered into the meniscus tissue’s inner, intermediate, and outer zones, respectively (**Figure 6a-b**). Alternatively, for more complex injuries, such as radial tears that begin in the inner meniscus and extend to the peripheral edge, the stiffness tunable DEM-based material system could be coupled with advanced fabrication techniques such as 3D bioprinting (**Figure 6a, c**). Employing our stiffness-tunable FADEM-based materials (‘soft’, ‘soft/stiff’, and ‘stiff’ FADEM), we crafted a hybrid hydrogel containing biomaterial and mechanical gradients. These gradients were produced using a multi-head 3D bioprinting system to mimic the zonal properties of the tissue (**Figure 6c**). FE-SEM analysis verified the retention of fibrous structures after printing (**Figure 6e-g**). Additionally, distinct zonal-dependent cell morphologies were observed when MSCs were cultured on the 3D printed construct, with smaller and rounder cells on the ‘soft’ hydrogel region (**Figure 6h**) and larger and elongated cells on the ‘stiff’ region (**Figure 6j**), with a mix of cell populations in the ‘soft/stiff’ region (**Figure 6i**). This highlights the capability of the system to effectively emulate the cell morphology of native meniscus tissue by adjusting hydrogel properties.

To further improve the printability of DEM-based materials and the integrity of the printed structures, we introduced a rapid curing reaction with the addition of ruthenium (Ru)/sodium persulfate (SPS) (**Figure 6k**). The Ru/SPS visible photoredox system interacts with tyrosine proteins prevalent in DEM, encouraging dityrosine bond formation for rapid cross-linking.[30] Through a triple crosslinking process utilizing heat, UV, and visible light during 3D bioprinting, we created 3D-printed hybrid structures with interconnected pores using ‘soft’ and ‘stiff’ FADEM-based hydrogels (**Figure 6k**). This improved structural fidelity not only benefits tissue engineering but also enhances the material’s physical and biochemical characteristics. The introduction of pores may support proper cell infiltration and differentiation based on the meniscus zone, potentially fostering the development of heterogeneous tissue structures that more closely resemble the native meniscus.

## Discussion

Current meniscus tissue engineering strategies are limited by their inability to replicate the complex zonal-dependent properties of meniscal tissue.^[1a]^ For instance, currently used biomaterials based on collagen and synthetic scaffolds do not mimic these zonal characteristics, leading to suboptimal healing and functional outcomes.^[1b]^ Similarly, traditional surgical interventions like suturing and arthroscopic partial meniscectomy (APM) fail to address the meniscus’s biochemical and structural heterogeneity in the meniscus tissue.[2]

To address these limitations, we introduced a novel approach for meniscus repair through the development of a tunable DEM-based hydrogel system, designed to approximate the zonal heterogeneity in the meniscus tissue. The goal in this study was to engineer a biomimetic material capable of emulating the distinct biochemical and mechanical profiles of the meniscus zones, thereby enhancing cell compatibility, guiding accurate zone-dependent cell differentiation, and improving healing outcomes. We successfully synthesized age-dependent DEM hydrogels (i.e., FDEM and ADEM) that reflect these zonal properties. These hydrogels influenced meniscus cell behavior, based on the age of the donor tissue used to prepare the DEM. For example, FDEM hydrogels favored chondrogenic gene expression, while ADEM hydrogels promoted fibrous gene expression. YAP localization studies further indicated differential mechano-activation on these two substrates. Proteomic analyses corroborated these findings, revealing a higher abundance of cell adhesion and fibrogenesis proteins in ADEM. Notably, the exclusive presence of fibronectin in ADEM, a pivotal protein for cell adhesion and proliferation, alongside its interaction with structural proteins like collagen, underscores the unique properties of the ADEM system in promoting cellular attachment and proliferation.[31] Additionally, blending different DEM types (i.e., FADEM) offers the opportunity to generate a spectrum of biochemical and mechanical stimuli. Moreover, unlike hydrogels requiring chemical additives (e.g., TGF-β3 or CTGF), our DEM hydrogels retain their age-specific properties post-decellularization, potentially offering a more advanced replication of the native tissue milieu and delivering biomechanical and biochemical signals that naturally modulate cell behavior. Indeed, the explant assays have confirmed that DEM hydrogels can recruit surrounding native tissue. Collectively, these findings demonstrate the unique capacity of DEM hydrogels to emulate the native tissue environment and provide tailored biomechanical and biochemical cues to direct cellular responses and, consequently, the success of regenerative therapies.

Despite their inherent advantages, DEM hydrogels alone may not fully capture the specific mechanical properties essential for recapitulating the region-specific properties of the meniscus. To overcome this, combined the age-dependent DEM with methacrylate hyaluronic acid (MeHA), a biomaterial known for its tunable mechanical properties.[32] This combination harnesses the biological benefits of DEM while providing further mechanical tunability via MeHA crosslinking, allowing for a more refined calibration of hydrogel stiffness. In addition, the decellularization process typically leads to a reduction in GAG content,[33] which is vital for biomechanical integrity in the meniscus. The addition of MeHA may compensate for this loss, replenishing hydration and mechanical attributes of the hydrogel. The MeHA-modified DEM hydrogels potentially create an optimized mechanical and biochemical milieu that promotes targeted cell proliferation and differentiation, mimicking the unique characteristics of various regions within the meniscus. Indeed, DEM-based ‘stiff’ MeHA hydrogels exhibit increased cell attachment and YAP nuclear localization, with FDEM-based ‘stiff’ MeHA hydrogels surpassing the effects observed with ADEM on its own. These findings imply that the DEM-MeHA hydrogel system not only facilitates mechanical and biochemical composition restoration but also regulates cellular activity. Consequently, DEM-MeHA hydrogels may present a more sophisticated and customized solution for meniscus repair.

In addition to its role in mechanical tuning, MeHA also contributes to improve the injectability of the DEM-based hydrogel, a critical feature for minimally invasive meniscus repair techniques.[34] This enhanced injectability could enable simpler biomaterial delivery and better *in situ* integration. MeHA also augments the bioprintability of the hydrogel, which is advantageous for 3D bioprinting applications, crucial for fabricating heterogeneous and anatomically precise scaffolds. The injectable and 3D-printable nature of the DEM-MeHA composite hydrogel broadens its clinical applicability, facilitating minimally invasive delivery and personalized scaffold production, thus improving the practicality and success rate of meniscus repair interventions. This study comprehensively evaluated the DEM-based hydrogel system through *in vitro*, *ex vivo*, and *in vivo* tests using a small animal model. However, further assessment in a more clinically relevant large animal meniscus injury model is still needed to validate its effectiveness for meniscus tissue repair following injection and transplantation of 3D-printed constructs. Additionally, while the current approach focuses on an acellular hydrogel system to enhance endogenous cell recruitment and promote tissue repair, future research should explore the synergistic effects of integrating cell-embedded hydrogels or soluble growth factors with injectable delivery and cell-printing technologies, offering a promising avenue for enhancing regenerative outcomes.

Taken together, our tunable DEM-based hydrogel system advances the field by providing a flexible approach to craft tailored microenvironments for tissue regeneration. The precise control over hydrogel stiffness and the ability to modulate cellular response based on both composition and stiffness enables targeted regenerative therapies, that may improve outcomes for meniscus repair and beyond. This study represents a significant advancement in tissue engineering, offering a promising platform for developing targeted regenerative therapies that could lead to improved clinical outcomes and a deeper understanding of mechanobiological influences in musculoskeletal repair. Furthermore, the adaptability of our hydrogel systems positions them as a formidable tool for a broad spectrum of biomedical applications, ranging from *in vitro* disease modeling to *in vivo* regenerative treatments.

## Materials and Methods

### Preparation of Age-Dependent Meniscus Decellularized ECM (DEM) Powder

Bovine fetal (3^rd^ trimester) and adult (~30-month-old) menisci for decellularization were purchased (Animal Technologies, Inc., USA) and meniscus tissues were decellularized following a previously reported protocol with minor adjustments.[30, 35] Briefly, the menisci were chopped into 2 mm^3^ cubes and stirred in deionized water for 4 hours to remove blood. Next, the tissues were decellularized in a solution of 0.3% (w/v) sodium dodecyl sulfate (Invitrogen, USA) in phosphate buffered saline (PBS; Growcells, USA) for 24 hours, followed by treatment with 3% (v/v) Triton-X100 (Sigma Aldrich; USA) in PBS for 24 hours, and 7.5 U/ml Deoxyribonuclease (Sigma Aldrich, USA) in PBS for 24 hours. To sterilize the treated tissues, tissues were rinsed with 4% ethyl alcohol for 4 hours. Residual chemicals were removed by rinsing the treated tissues with PBS between each treatment step. After lyophilization, the tissues were crushed into powder using a liquid nitrogen cryomill (Rate: 14 cycle per second) to yield fetal and adult meniscus dECM powders.

### Fabrication of Age-Dependent Meniscus DEM Hydrogels

To prepare DEM-based hydrogels, the fabricated fetal and adult meniscus DEM powders were each digested in a 0.5 M acetic acid solution (Sigma-Aldrich, USA) with pepsin (Sigma-Aldrich, USA) at room temperature for 72 hours. After filtration through a 40 μm pore mesh, the digested solution was neutralized using a 10 N NaOH solution (Sigma-Aldrich, USA). The DEM pre-gel was prepared by adding 10× PBS and sterilized water into the neutralized DEM solution. Fetal DEM (FDEM) and adult DEM (ADEM) hydrogels were fabricated by cross-linking the prepared pre-gel at 37℃ for 30 mins. To create a composite group comprised of both the fetal and adult DEM, fetal and adult DEM (FADEM), the prepared FDEM and ADEM pre-gels were mixed in a 1:1 ratio. A 1.5 wt% concentration of FADEM was used for experiments.[33] To examine fibrous structures in DEM, the fabricated DEM gels were dried at room temperature, coated with platinum using a sputter coater, and imaged using a FEI Quanta FEG 250 scanning electron microscope (SEM; ThermoFisher, USA) at magnification of 5,000×. The distribution of fiber diameters was analyzed using ImageJ (National Institutes of Health, USA).

### Protein Identification by Mass Spectrometry (LC-MS/MS)

To analysis protein components in the fabricated Fetal and Adult DEM, proteomic analysis was performed. To prepare samples, the fetal and adult bovine menisci with four different donors were decellularized in the same protocol as prementioned. The DEM powder was fabricated through the lyophilization and cryomill procedures and used for the proteomic analysis. A mass spectrometry (LC-MS/MS) was performed as described in Supplementary information. Only proteins detected in two or more of the four different donors were processed for analysis. Differential level of proteins was identified with criteria of p-value < 0.05 and absolute log_2_ (Fold change) > 1.0 for volcano plot. Gene symbols corresponding to the identified proteins were input into the STRING web tool (version 10.0, http://string.embl.de, String Consortium 2020) to obtain data on protein-protein networks.[36] The acquired data were then imported into Cytoscape (Version 3.10.1) for visualization and annotation.[37] Gene ontology (GO) analysis was conducted using the STRING web tool, which identified biological process (BP), cellular component (CC), and molecular function (MF) terms with a corrected p-value < 0.05.

### Fabrication and Characterization of Stiffness Tunable DEM-based MeHA Hydrogels

Sodium hyaluronic acid (HA; Lifecore Biomedical, USA) was methacrylated as previously described.[32, 38] Briefly, methacrylate HA (MeHA) was obtained by methacrylate esterification with the hydroxyl group of the 68 kDa sodium HA. The degree of the methacrylation was controlled by adjusting the amount of methacrylic anhydride (Sigma Aldrich, USA), with target methacrylation degrees of ~30% (‘soft’) and 100 % (‘stiff’). After dialysis at room temperature, stiffness-modulated MeHA macromers were isolated by freezing and lyophilizing. Lyophilized polymers were dissolved in deuterium oxide (Sigma Aldrich, USA) at a concentration of 10 mg/mL and analyzed using ^1^H-NMR (Bruker NEO400, USA) to determine the degree of modification.[39]

To fabricate DEM-based MeHA hydrogels with different stiffness, age-dependent DEM (FDEM or ADEM) and the stiffness-tuned MeHA system (soft or stiff) was blended to a final concentration of 1.5 wt% and 1.0 wt% final concentrations, respectively. In another group, soft and stiff MeHA were mixed at a 1:1 ratio. Thiolated fluorescein peptide (GenScript, USA, 2.0-2.5%) was incorporated during crosslinking to observe the homogeneous blending of the FITC-Soft/Stiff MeHA and DEM. The thiol groups from the fluorescein peptide and the methacrylate groups from the MeHA macromers were conjugated via Michael addition reactions.[40] After dialysis, the fluorescein peptide-conjugated MeHA macromers were isolated through freezing and lyophilization. To assess the homogeneity of these macromers, a confocal microscope (TCS SP8 STED, Leica, Germany) was employed. The macromers were scanned over an area of 1.5 mm × 1.5 mm to a depth of 200 μm (Z stack; 10× magnification) to evaluate their distribution and uniformity.

A custom mechanical testing device was used to evaluate compressive moduli of the fabricated hydrogels.[41] For this, the hydrogel samples were prepared in cylindrical molds measuring 5 mm diameter and 2 mm thickness. Samples were equilibrated in creep under a static load of 0.1 g for 5 mins. After creep, samples were subjected to 50 % strain applied at 0.5%/s. The compressive modulus was determined from the stress (minus tare stress) normalized to the applied strain in the linear region.

### Cell Isolation and *In Vitro* Culture

Cell isolation for in-vitro testing was performed as previously reported.[42] Briefly, juvenile bovine knee joints (2-3 months old; Research 87, USA) were acquired. Bone marrow derived mesenchymal stem cells (MSCs) were isolated from the femur and tibia, and meniscal fibrochondrocytes (MFCs) were isolated from the meniscus. Cells were cultured in basal growth media [BM; Dulbecco’s modified Eagle’s medium (DMEM; Gibco, USA) supplemented with 10% fetal bovine serum (R&D systems, USA) and 1% Penicillin Streptomycin (Gibco, USA)]. Cells at passage 2 were used for in-vitro experiments. To prepare ADEM-conditioned media (ACM), ADEM was added to fresh BM and incubated at 37°C with shaking for 5 days. Concurrently, fresh BM was also incubated under the same conditions with ACM, serving as the control group (Ctrl). After filtration through a 40 μm pore mesh, the ACM with a 3.3% v/v concentration was used for the dissolution test *in vitro*, having confirmed its low cytotoxicity via a Live/Dead assay. The DEM system was coated onto a 24-well tissue culture plate and crosslinked in a CO2 incubator for 30 minutes. For the DEM-based MeHA system, an additional crosslinking step was performed by exposing it to 15-20 mW/cm² UV light (Omnicure S2000-XLA, Lumen Dynamics, Canada) for 30 minutes. Following crosslinking, cells were seeded on the surface of the coated DEM or DEM-based MeHA systems and cultured in either BM or ACM.

### Metabolic and Cytotoxicity Assays

The DEM-based MeHA hydrogel was added to a 24-well plate to completely cover the bottom surface. Cells were then seeded at a density of 1 × 10^5^ cells per well (n=5/group). Proliferation of seeded cells was determined using a Cell Counting Kit-8 assay (CCK-8; Dojindo, Japan). After 1, 3, and 7 days, medium containing 10% CCK-8 agent was added to each well and incubated at 37 ℃ for 4 hours. The absorbance of each well was measured at 450 nm using a Synergy H1 microplate reader (BioTek, USA). Cell viability was assessed using the Live/dead kit (Invitrogen, USA). Live cells were stained by Calcein AM and dead cells were stained by Ethidium homodimer-1. After imaging, quantification was performed in ImageJ (n=5).

### Analysis of Cell Morphology and YAP Nuclear Localization

Immunofluorescence staining was performed to measure cell morphology and YAP (Yes-associated protein) in cells cultured on hydrogels. Cells were fixed at day 3 with 4% paraformaldehyde (PFA). To examine cell morphology, cells were permeabilized with Triton X-100 and stained with Phalloidin-Alexa488 (Invitrogen, USA) to visualize actin. Nuclei were labeled with ProLong^TM^ Gold antifade reagent with DAPI (Invitrogen, USA). For YAP staining, cells were incubated with anti-YAP antibody (Santa Cruz Biotechnology, USA) followed by Alexa Fluor546-goat anti-mouse IgG (Life technologies, USA). Images were obtained using a widefield fluorescent microscope (Leica, Germany). Cell morphology and YAP nuclear localization were analyzed with ImageJ

### Gene Expression Analysis

Messenger RNA was extracted from the MSCs or MFCs cultured on hydrogels using the TRIzol/chloroform method[41]. The concentration of the total RNA was determined using a spectrophotometer (Nano-drop Technologies, USA)^[38a]^. Subsequently, cDNA was synthesized using PhotoScript II First-Strand cDNA Synthesis Kit (New England BioLabs, USA). Reverse transcription-polymerase chain reaction (RT-PCR) was performed on a QuantStudio 6 Pro (Applied Biosystems, USA) with Fast SYBR Green Master Mix (Applied Biosystems, USA). The expression of collagen type I alpha 2 chain (COL1A2), Collagen Type-2 (COL2), Collagen Type-3 (COL3), Connective tissue growth factor (CTGF), Aggrecan (ACAN), SRY-box Transcription Factor 9 (SOX9), Transforming growth factor (TGF), Matrix metallopeptidase 1 (MMP1), and a disintegrin and metalloproteinase with thrombospondin motifs 4 (ADAMTS4) were determined. Expression levels were normalized to the house-keeping gene Glyceraldehyde 3-phosphate dehydrogenase (GAPDH). Primer sequences are provided in **Table 1**.

### RNA sequencing

RNA sequencing was carried out with the MSCs cultured on FADEM-based ‘Soft’, ‘Stiff’ hydrogels. Each type of hydrogel was coated on the bottom surface of 24-well tissue culture plates and cross-linked under UV and 37°C. Passage 2 (P.2) cells were seeded in the 24-well tissue culture dishes at a density of 1×10^5^ cells per well. The cells were cultured in basal growth media for 7 days. Total RNA was extracted from the cells on the hydrogel surface using the TRIzol/chloroform method.[41] RNA-Seq library preparation (including rRNA depletion) and sequencing (150 bp paired-end) were performed at Genewiz (South Plainfield, New Jersey). Sequence reads were processed using Trimmomatic v.0.36 to trim potential adapter sequences and low-quality nucleotides. The processed reads were aligned to the Bos taurus reference genome (available on Ensembl) with the STAR aligner v.2.5.2b. Unique gene hit counts were determined using featureCounts from the Subread package v.1.5.2. Differential expression analysis between the experimental conditions, FADEM-based Soft MeHA and FADEM-based Stiff MeHA, was conducted using the DESeq2 package in R. Global transcriptional changes across the groups were visualized with GraphPad Prism. The DESeq2 Likelihood Ratio Test (LRT) was employed to assess differential expression between the contrasts of interest, with a stringent significance threshold of p < 0.05 to ensure a high level of confidence in rejecting the null hypothesis. Differentially expressed genes (DEGs) were identified using the DESeq2 package with criteria of Benjamini-Hochberg adjusted p-value < 0.05 and absolute log2 (fold change) > 1.0. The R pheatmap package was utilized to illustrate the expression patterns, clusters, and distribution of significant genes from the LRT analysis. To achieve a deeper insight into the functional profiling of genes significant in the LRT analysis, a gene enrichment analysis was conducted to identify highly enriched biological processes. This analysis was performed using the g function on the gProfiler web server (https://biit.cs.ut.ee/gprofiler/gost). The significance threshold was set using the Benjamini-Hochberg FDR (false discovery rate), with significant Gene Ontology (GO) terms defined by an adjusted p-value of <0.05. For visualization, the significant GO terms were created using the ggplot2 package in R.

### Ex-vivo Evaluation of Age-Dependent Meniscus DEM Hydrogels in Meniscus Defects

Whole menisci were dissected from fresh juvenile bovine knees, and radial incisions were made. Using biopsy punches, a 3 mm diameter hole was created in the avascular zone of each cross-section, and the DEM-based hydrogel was injected into the holes.[43] (**Figures 2i).** The defect group, serving as a control, was left unfilled. The tissues were cultured for 3 weeks in basal growth media. For histological analysis, tissues were fixed in 4% paraformaldehyde, embedded in Cryoprep frozen section embedding medium, and sectioned at a thickness of 8 µm. Hematoxylin and eosin (H&E) staining was carried out to evaluate the DEM matrix preservation and cell infiltration during the culture period. The number of nuclei present in the H&E images was counted using ImageJ. Picrosirius red (PSR) staining was performed to visualize collagen. Additional tissues with injected DEM gel were fixed in 4% paraformaldehyde, infiltrated with Citrisolv, embedded in paraffin, and sectioned at 3 weeks. H&E staining was conducted to confirm cell recruitment into the injected DEM gel. The nuclear area on the H&E images was measured in ImageJ, and Picrosirius red (PSR) staining was performed to visualize collagen.

### In-vivo Explant Evaluation of FADEM-based Hydrogels in Meniscus Defects using mouse subcutaneous model

All animal studies were conducted under the animal research protocol (No. 22-268) approved by the Committee of the Use of Live Animals in Teaching and Research (CULATR) at the University of Hong Kong. The studies adhered to the Animals (Control of Experiments) Ordinance (Hong Kong) and guidelines from the Centre for Comparative Medical Research (CCMR), Li Ka Shing Faculty of Medicine, The University of Hong Kong. Male severe combined immunodeficient (Severe Combined Immunodeficiency; CCMR) mice (6-8 weeks old) were used for transplantation experiments. The mice were obtained from the Experimental Animal Center of the University, with access to food and water ad libitum, and were maintained under pathogen-free conditions. All procedures were performed under anesthesia using intraperitoneal injections of ketamine/xylazine mixtures [100 mg/kg ketamine (Cat. No. 013004, AlfaMedic Ltd.,) + 10 mg/kg xylazine (Cat. No. 013006, AlfaMedic Ltd.)]. Anesthesia was confirmed by checking body reflexes, and eye ointment (Duratears® ointment, Cat. No. 05686, Alcon, Fort Worth, TX, USA) was applied to prevent blindness.

The biocompatibility and stability of the FADEM-based hydrogel were evaluated by injecting ‘soft’ or ‘stiff’ FADEM-based MeHA hydrogels into juvenile bovine meniscus tissue explants, which were prepared by punching and cutting fresh bovine meniscus tissue into discs of 2 mm in diameter and 5 mm in height. After hydrogel injection, the samples were cross-linked using UV light and incubated with CO_2_ for 30 minutes, followed by a 5-day preculture period before subcutaneous implantation into athymic mice (**Supplementary Figure S9a**). The subcutaneous implantation sites on the mice were decontaminated with alternating applications of 70% alcohol and povidone-iodine (Betadine®) using sterile cotton swabs. The hydrogel-laden bovine meniscus tissue explants were then implanted into the left and right backs of the mice. After suture, the mice were placed in an intensive care unit (ICU) until they fully recovered. Finally, they were returned to their original housing location. The three weeks post-implantation, the mice were euthanized by an overdose of Pentobarbital (250 mg/kg) (Dorminal®, Cat. No. 013003, AlfaMedic Ltd.) administered intraperitoneally. The implant areas were harvested and immediately fixed in 10% neutral-buffered formalin (NBF) for 24 hours for further histological analysis. The samples underwent a series of ethanol, xylene-ethanol, and xylene treatments (1 hour each, then were embedded in paraffin and sectioned at 10 μm thickness using a rotary microtome (Cat. No. RM2155, Leica Microsystems, Wetzlar, Germany).

Series of sectioned slides were deparaffinized, rehydrated, and stained with hematoxylin (Cat. No. SH4777, Harris Hematoxylin, Cancer Diagnostic Inc., USA) and eosin (Cat. No. CS701, Dako, Denmark) (H&E) and Picrosirius red (Cat. No. PH1098, Scientific Phygene^®^). All staining operations were performed in accordance with the company product’s instructions and guidelines. After dried and mounted through xylene-based solution, the slides were then observed under an optical microscopy (Eclipse LV100POL, Nikon, Japan) with a digital video camera (Digital Sight DS-Ri1, Nikon, Japan).

### Extrusion-based 3D Printing

FADEM-based hydrogels were printed using a BIO X 3D bioprinter (Cellink, Sweden). To prepare the FADEM-based MeHA hydrogel, FDEM and ADEM hydrogels were neutralized with 10 M NaOH. A blend of FADEM blended with ‘soft,’ ‘soft/stiff,’ or ‘stiff’ MeHA was created, and the final concentration was designed to be 1.5 wt% FADEM-1.0 wt% MeHA. Then, they were loaded into syringes and stored at 4℃. Extrusion was performed using 22 G nozzles with pneumatic pressures ranging from 10 to 30 kPa. The formed hydrogels were exposed to 18-24 mW/cm² UV light (365 nm wavelength) during the extrusion process. To distinguish the printed materials, Rhodamine B and Fluorescein disodium salt (AquaPhoenix Scientific, USA) were added before the 3D printing process. To enhance the printability of the DEM-based hydrogel systems, Ruthenium/sodium persulfate (RS, 1/10 × 10⁻³ M R/S) was added to the DEM-MeHA hydrogels. The same printing conditions were maintained, with visible light (>405-450 nm wavelength) at 10-30 mW/cm² applied simultaneously during material extrusion.[30]

### Statistical Analysis

All data are presented as the mean ± standard deviation. Statistical analysis was performed using one-way analysis of variance (ANOVA) with Tukey’s *post hoc* testing with a 95% confidence interval via GraphPad Prism version 9 software (San Diego, USA). A p-value less than 0.05 was considered statistically significant.

## Acknowledgments

This research was supported by the National Institutes of Health (K01 AR07787, R21 R077700, P30 AR069619, R01 AR056624), National Science Foundation (CMMI 1548571), Department of Veterans Affairs (CReATE Motion Center, 1I50RX004845-01) and the Korea Health Technology R&D Project through the Korea Health Industry Development Institute (KHIDI), funded by the Ministry of Health and Welfare, Republic of Korea (HI19C1095).

## Author Contributions

S.-H.L., Z.L., E.Y.Z., D.H.K., Z.H., S.J.L., H.-W.K., J.A.B., R.L.M. and S.C.H. designed the studies. SHL, ZL, EYZ, DHK, and ZH performed the experiments. S.-H.L., Z.L., E.Y.Z., D.H.K., Z.H., S.J.L., H.-W.K., J.A.B., R.L.M. and S.C.H. analyzed and interpreted the data. S.-H.L., Z.H., S.J.L., R.L.M. and S.C.H. drafted the manuscript, and all authors edited the final submission.

## Supplementary Information

### Supplementary Materials and Methods

#### Proteomic Analysis by Mass Spectrometry (LC-MS/MS)

##### In-solution digestion

The lyophilized, decellularized tissue underwent digestion following a previously outlined procedure.[1] Initially, samples were solubilized in 8M urea, followed by reduction, alkylation, and subsequent dilution to 2M urea. PNGaseF treatment was then applied to remove N-linked glycans. Ammonium bicarbonate was introduced to achieve a final concentration of 100 mM, initiating proteolysis with Lys-C at 37°C for 2 hours. Subsequently, trypsin was added, and the samples were incubated overnight at 37°C. An additional trypsin incubation for 2 hours followed. The resulting samples were acidified, desalted, and dried through vacuum centrifugation. Prior to LC-MSMS analysis, all peptides were solubilized in 0.1% TFA supplemented with iRT peptides (Biognosys AG, iRT).

##### Mass spectrometry data acquisition

Samples were randomized and subjected to analysis on a Exploris 480 mass spectrometer (Thermo Fisher Scientific, San Jose, CA) coupled with an Ultimate 3000 nano UPLC system and an EasySpray source. Data were acquired using data independent acquisition (DIA). Tryptic digests were supplemented with iRT standards (Biognosys) and separated by reverse-phase (RP)-HPLC using a nanocapillary column (75 μm ID × 50 cm, 2 μm PepMap RSLC C18 column) at 50°C. The mobile phases consisted of 0.1% formic acid (mobile phase A) and 0.1% formic acid/acetonitrile (mobile phase B). Peptides were eluted into the mass spectrometer at a flow rate of 300 nL/min, with each RP-LC run encompassing a 180-minute gradient from 1 to 5% B over 15 minutes, followed by 5-45% B.

##### Data-independent acquisition (DIA) setting

DIA mass spectrometer settings were configured as follows: one full MS scan at 120,000 resolution, covering a scan range of 350-1200 m/z with a normalized automatic gain control (AGC) target of 300% and automatic maximum inject time. This was followed by variable isolation windows for DIA, MS2 scans at 30,000 resolution, a normalized AGC target of 1000%, and automatic injection time. The default charge state was set to 3, the first mass was fixed at 250 m/z, and the normalized collision energy for each window was established at 27.

##### QA/QC and system suitability

The performance of the Exploris 480 instrument was assessed using QuiC software (Biognosys; Schlieren, Switzerland) for analyzing the spiked-in iRT peptides. To ensure quality control, standard E. coli protein digest injections were interspersed between samples (one injection after every four biological samples), and the data were collected in data-dependent acquisition (DDA) mode. Subsequently, the acquired DDA data were processed using MaxQuant,[2] and the results were visualized using the PTXQC package to monitor the instrument’s quality.[3]

##### Mass spectrometry raw data processing

The MS/MS raw files underwent processing in Spectronaut 15.7 (Biognosys AG).[4] For this analysis, we utilized a reference Bovine proteome from Uniprot, comprising 37,879 proteins (downloaded on 19052022), supplemented with a list of 245 common protein contaminants and iRT peptides. Trypsin was designated as the enzyme with allowance for two possible missed cleavages. Carbamidomethyl of cysteine was defined as a fixed modification, while protein N-terminal acetylation and oxidation of methionine were considered variable modifications. A false discovery rate limit of 1% was applied for peptide and protein identification, with the remaining search parameters set to their default values.

##### Proteomics data processing and statistical analysis

Proteomic data processing and statistical analysis were performed in Perseus.[5] The MS2 intensity values generated by Spectronaut were used to analyze the whole proteome data. The data were log2 transformed and normalized by subtracting the median for each sample. We filtered the data to have a complete value for a protein in at least one cohort. To compare proteomics data between groups, a t-test was employed to identify differentially expressed proteins, and volcano plots were generated in visualize the affected proteins while comparing different groups. Lists of differentially abundant proteins was then sorted based on the adj.P.value <0.05 and |log_2_FC| > 1, yielding a prioritized list for downstream bioinformatics analysis.

### Supplementary Figures and Legends

**Supplementary Figure S1.**
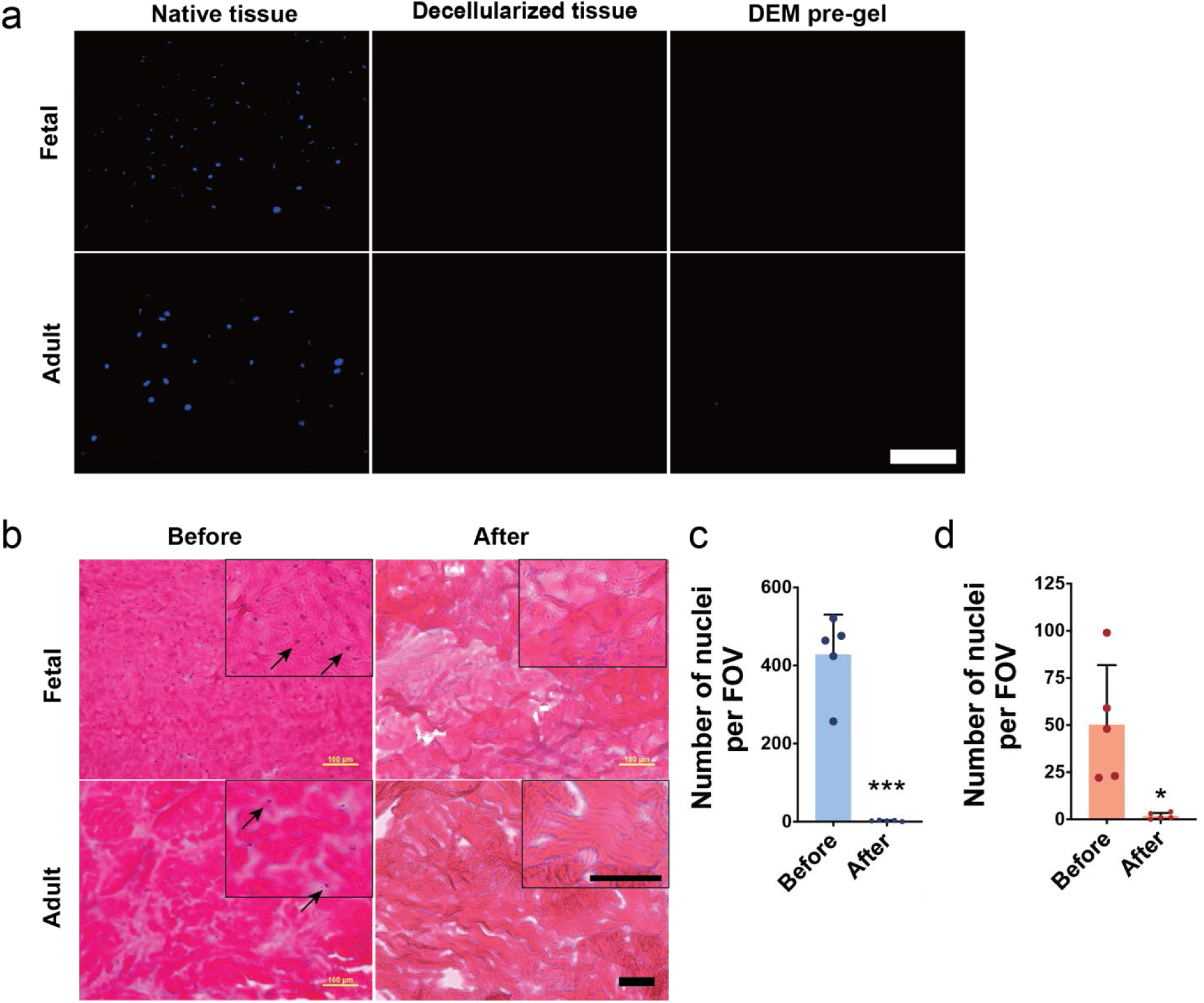
a) Representative images of DAPI stained tissue sections before and after decellularization and in the pre-gel formulation (scale bar: 100 µm). b) Representative images of H&E stained sections (arrows indicate nuclei; scale bar: 100 µm). Quantification of the number of nuclei per field of view (FOV) in c) Fetal and d) Adult tissue before and after decellularization (n=5; *p<0.05 vs. Before, ***p<0.001 vs. Before).

**Supplementary Figure S2.**
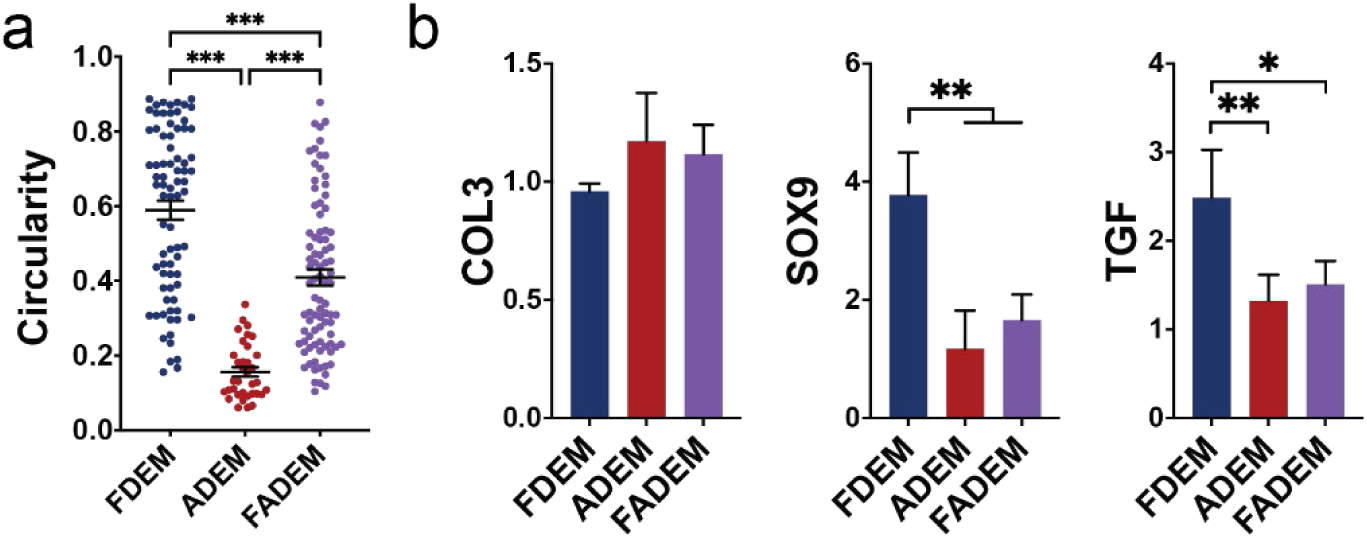
MSCs in in vitro culture: a) Circularity (n=35-87) on day 3, and b) gene expression on day 7 (n=3-4); *p<0.05, **p<v0.01.

**Supplementary Figure S3.**
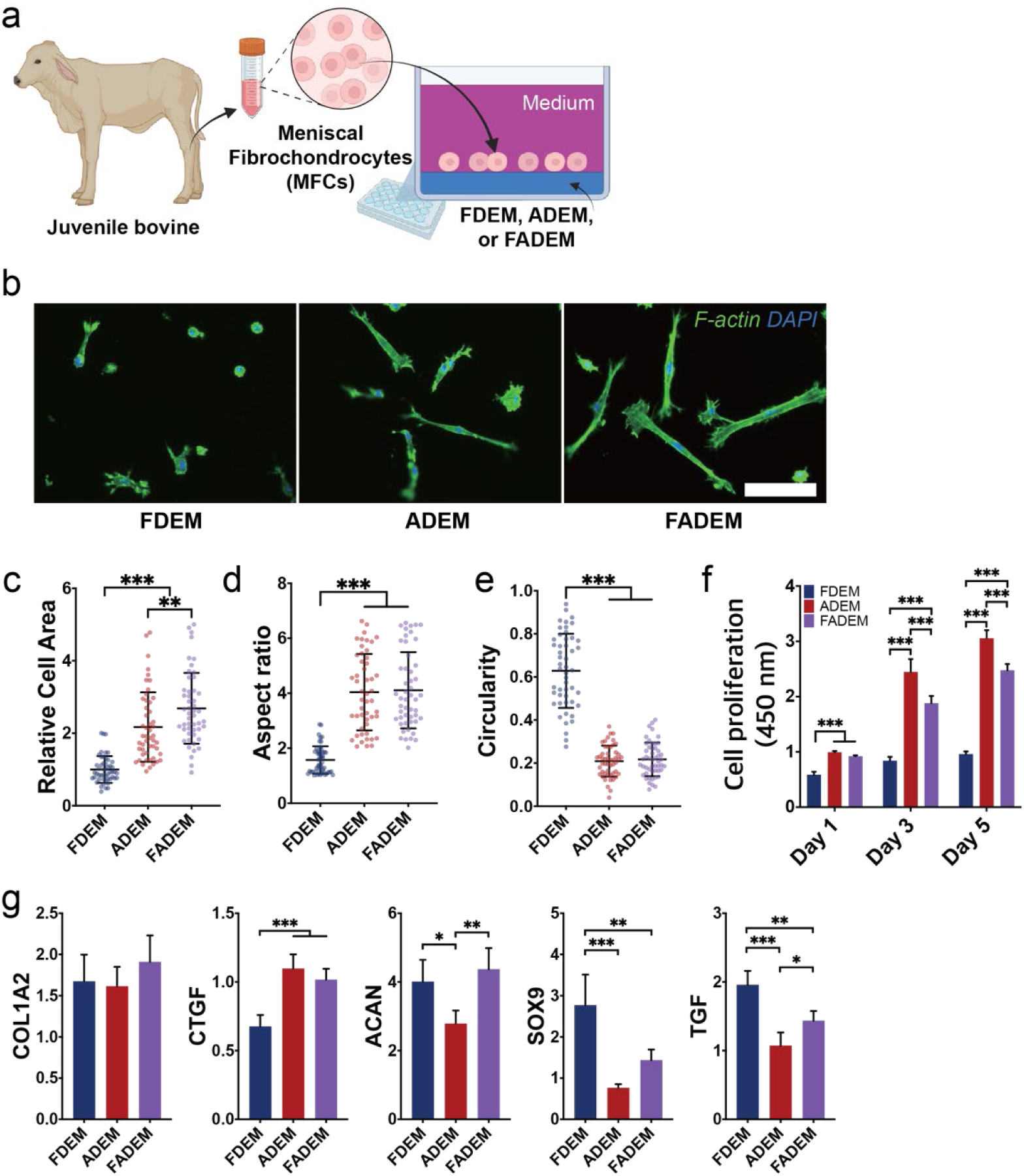
Meniscus fibrochondrocytes (MFCs) in in vitro culture: a) Illustration of in vitro experiment, b) Representative F-Actin images (Scale bar: 100 µm), c) relative cell area, d) aspect ratio, and e) circularity on Day 3 (n=50); **p<0.01, ***p<0.001). f) Cell proliferation (n=5) and g) gene expression on Day 5 (n=5); COL2 was not detected; **p<0.01, ***p<0.001.

**Supplementary Figure S4.**
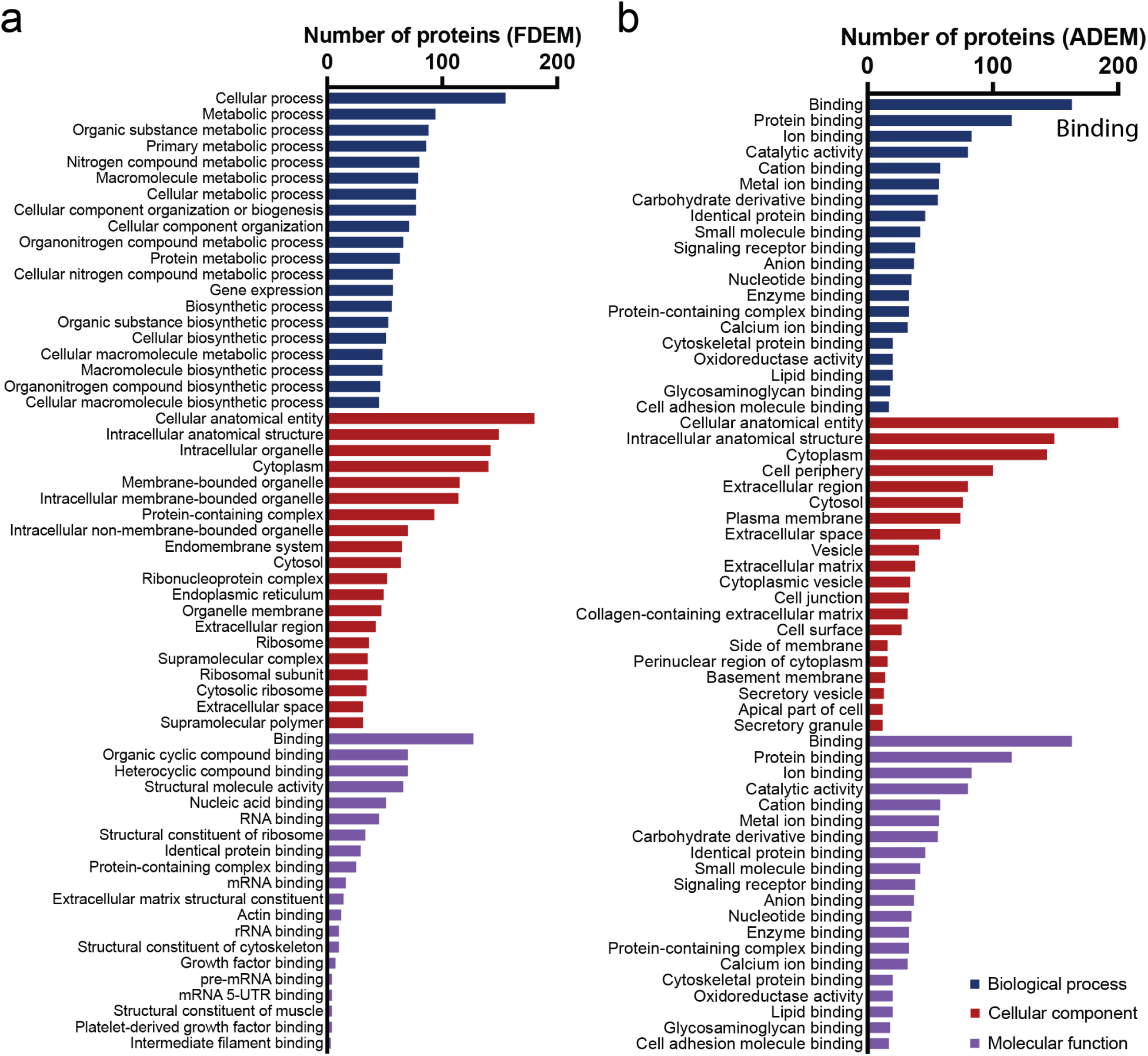
In-depth gene ontology (GO) analysis for both FDEM and ADEM, derived from four distinct donors. The analysis encompasses: a) FDEM: Showcasing the top 20 GO terms associated with Biological Process, Cellular Component, and Molecular Function, highlighting the cellular activities and interactions prevalent in FDEM, b) ADEM: Similarly, detailing the top 20 GO terms for ADEM, emphasizing the protein, enzyme, and ECM binding activities that are characteristic of this group.

**Supplementary Figure S5.**
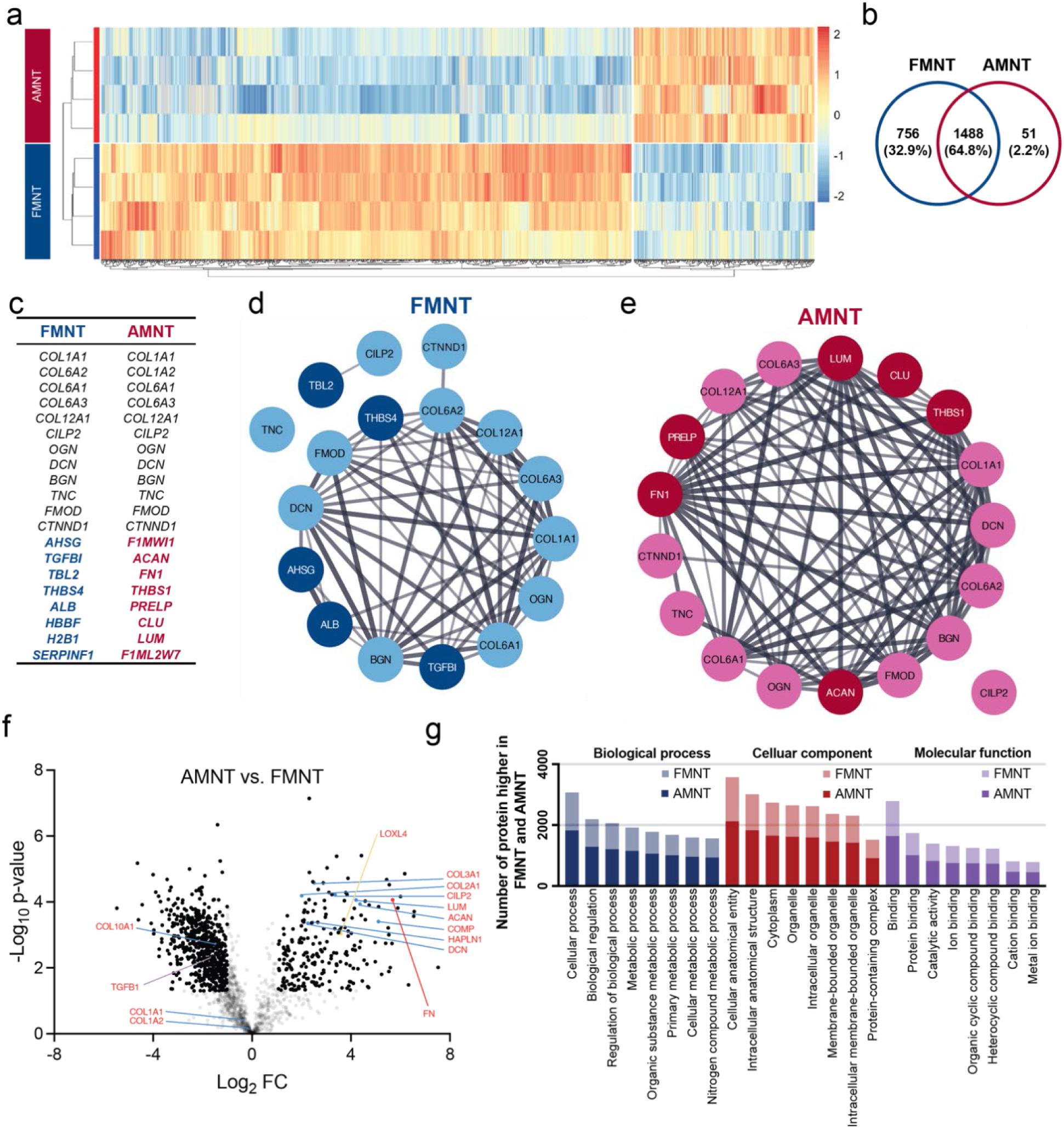
Proteomic landscape of native fetal meniscus tissue (FMNT) and native adult meniscus tissue (AMNT), with each group comprising four different donors. a) Heatmap and clusters demonstrating the protein detection patterns in FMNT and AMNT, indicating the proteomic distinctions between groups, b) Venn diagram illustrating the shared and unique proteins in FMNT and AMNT, c) Listing of top 20 most abundant proteins detected in both native tissues, d-e) Protein-Protein Interaction Networks for the top 20 proteins of FDEM (d) or ADEM (a) with line thickness representing the strength of data support for each interaction, f) Volcano Plot of AMNT (vs. FMNT) showing the differentially abundant proteins between AMNT and FMNT, with statistical significance and fold-change metrics, g) Gene Ontology Analysis indicating the top 8 GO categories for Biological Process, Cellular Component, and Molecular Function, with color-coding to differentiate the categories that vary between FMNT and AMNT.

**Supplementary Figure S6.**
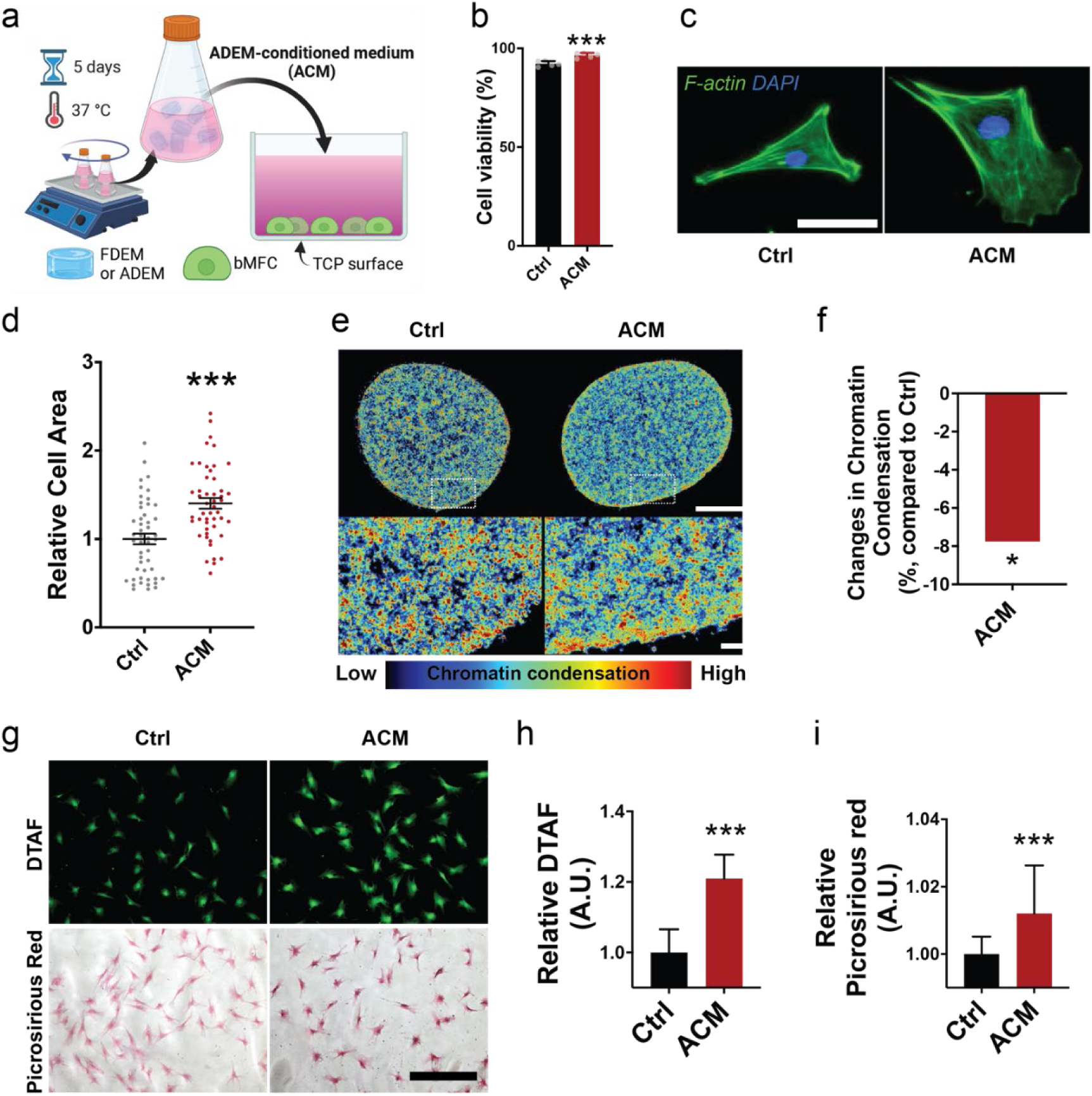
a) Schematic of experiment setup for the conditioned media assay, b) Cytotoxicity results after 5 days (n=5; ***: p<0.001), c) Representative F-actin images from day 3 (Scale bar: 50 μm), d) Quantified cell area (n=46-49; ***: p<0.001 vs. Ctrl), (e) Representative images showing chromatin condensation levels in MFCs [Scale bars: 1 μm (top) and 500 nm (bottom)], and f) Chromatin condensation quantification (*: p<0.05 vs. Ctrl). Protein staining: g) Representative images of DTAF staining and Picrosirius Red staining (Scale bar: 200 um) of MFCs. Relative intensity measurements of the DTAF (h) and PSR (i) staining intensity per cell (n=25; *: p<0.05 vs. Ctrl, ***: p<0.001 vs. Ctrl).

**Supplementary Figure S7.**
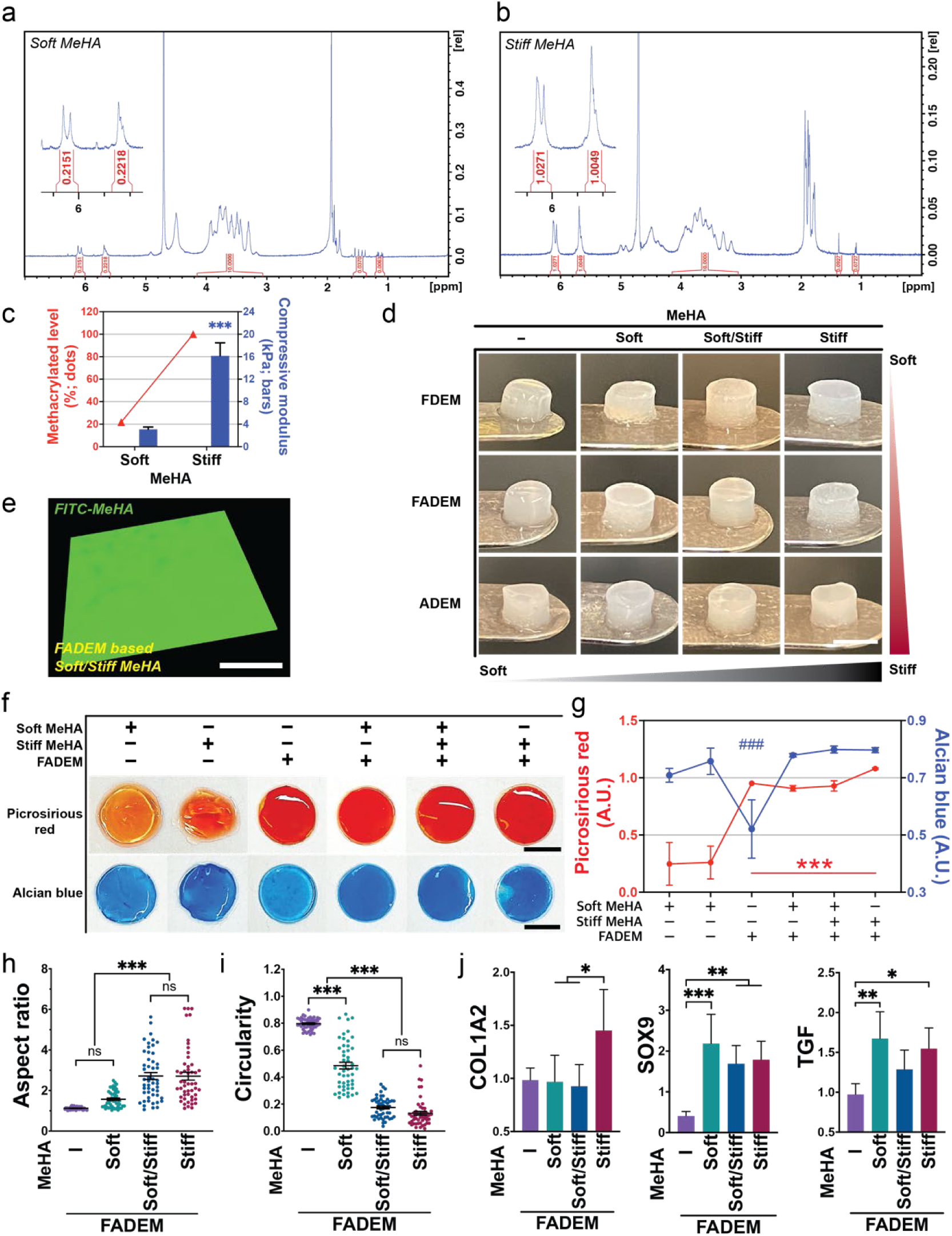
a) Degree of methacrylation analysis by ¹H-NMR: Soft MeHA. b) Stiff MeHA. c) % Methacrylation and Mechanical Properties: Comparison of % methacrylation and mechanical properties of MeHA hydrogels (n=5; ***p<0.001 vs. Stiff MeHA), d) Images of fabricated stiffness tunable HA/DEM hydrogels (scale bars: 3 mm). e) Fluorescence images showing homogeneity: ‘soft/stiff’ FADEM-based hydrogels (Green: ‘soft/stiff MeHA distribution; Scale bar: 400 µm), f) Collagen and proteoglycan staining: representative images of Picrosirius red (PSR) and Alcian blue (AB) staining, g) Quantification of PSR and AB staining intensity (n=5; ###: p<0.001 vs. whole groups in Alcian blue graph; ***: p<0.001 vs. Soft MeHA in PSR graph). Quantification of cell aspect ratio (h) and circularity (i) (n=50; ***p<0.001). j) Quantification of gene expression by MSCs on stiffness-tunable HA/DEM hydrogels (n=4-5; *p<0.05, **p<0.01, ***p<0.001).

**Supplementary Figure S8.**
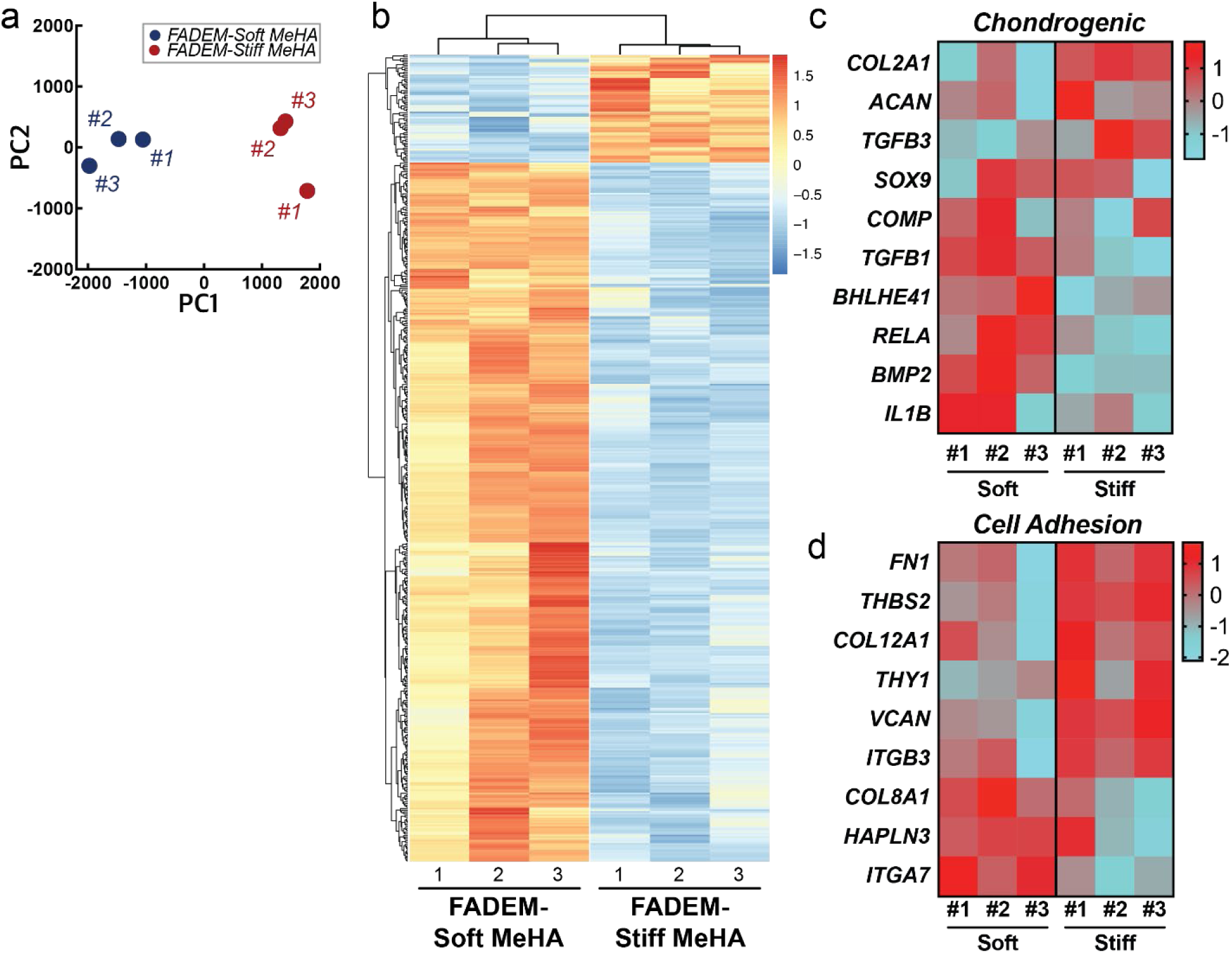
a) PCA plot clustering of samples, b) Heatmap showing expression patterns of significantly altered genes across different experimental conditions. Z-score heat maps of Chondrogenic (c) and cell adhesion (d) related gene expression (n=3/group).

**Supplementary Figure S9.**
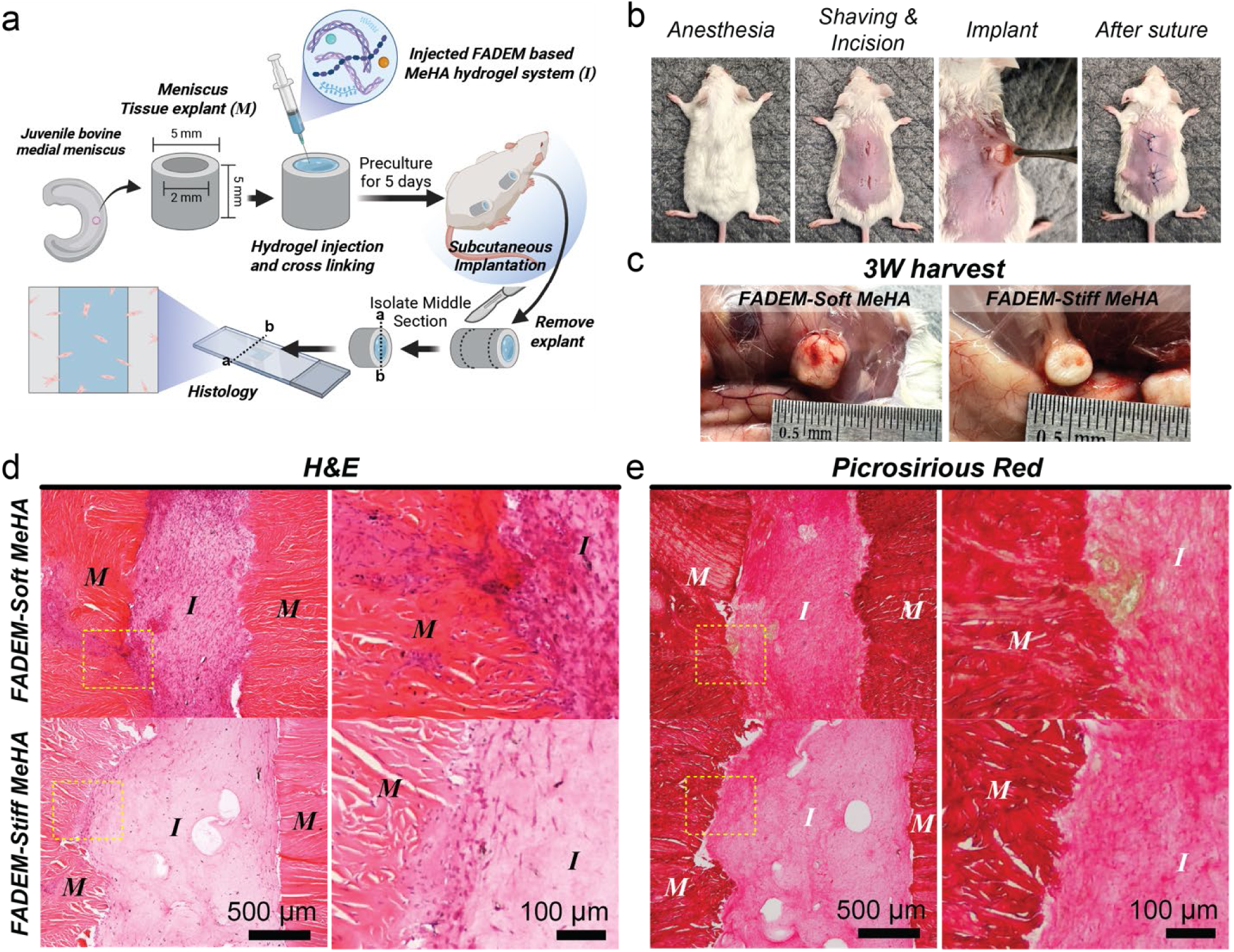
a) Schematic illustration of bovine tissue implant with injected FADEM-based tunable MeHA hydrogel, subcutaneous implantation in mice, and preparation for histological staining, b) Images showcasing the surgical procedure, c) Harvested samples at 3 weeks post-implantation. Histological Analysis (3 weeks post-implantation): d) Representative H&E-stained and e) Picrosirius red-stained sections (M: Meniscus tissue explant, I: Injected FADEM-based MeHA hydrogel system).

**Supplementary Table 1.**
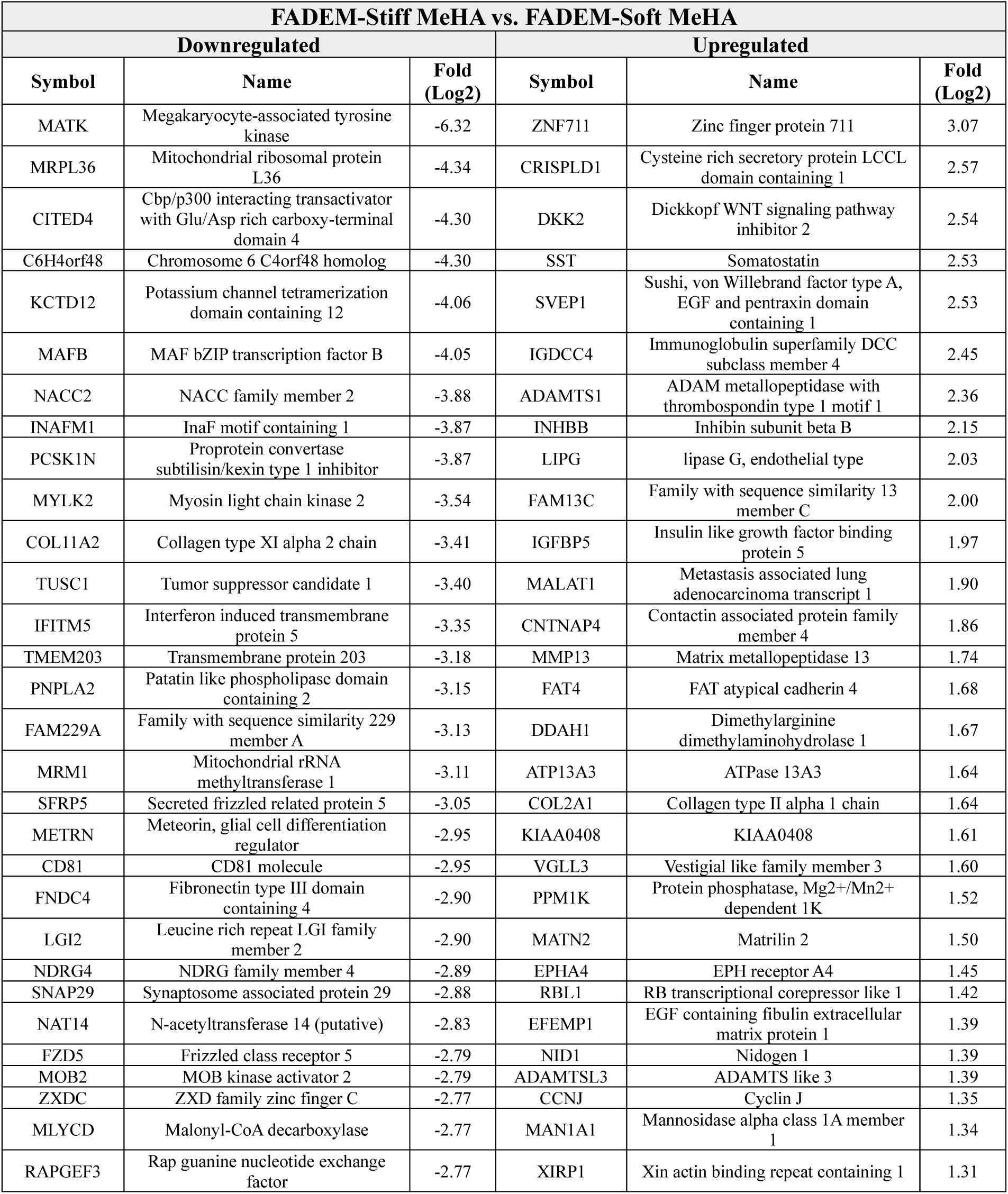
Top 30 up- or down-regulated genes.

**Supplementary Table 2.**
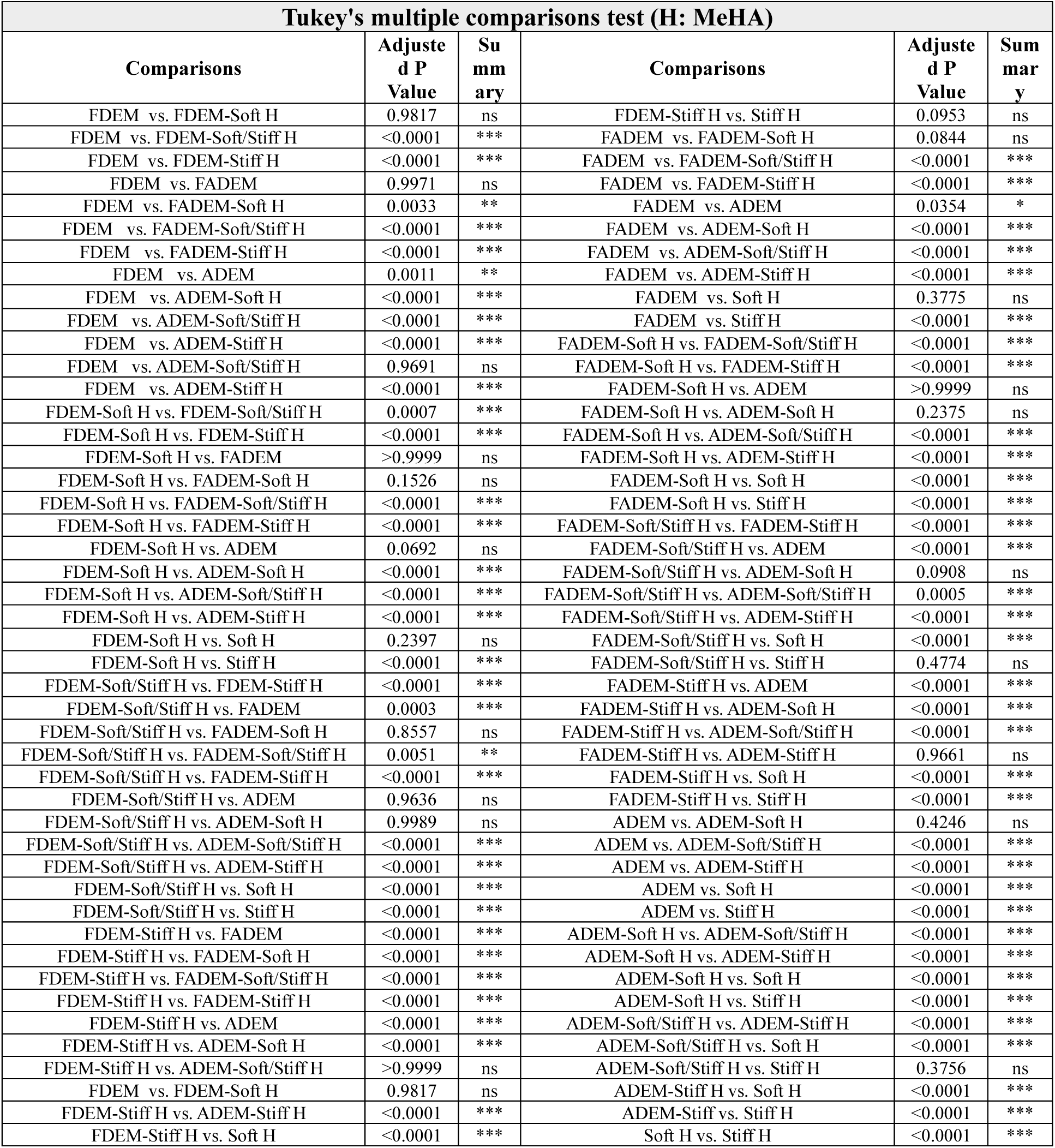
Tukey’s multiple comparisons test for stiffness of mechanical modulated DEM-based MeHA hydrogel system (Figure 5b; n=5; ns: not significant, *<0.05, **p<0.01, ***p<0.001)

## Supplementary References

[1] A. Naba, K. R. Clauser, R. O. Hynes, JoVE (Journal of Visualized Experiments) 2015, e53057.

[2] S. Tyanova, T. Temu, J. Cox, Nat. Protoc. 2016, 11, 2301.

[3] C. Bielow, G. Mastrobuoni, S. Kempa, J. Proteome Res. 2016, 15, 777.

[4] R. Bruderer, O. M. Bernhardt, T. Gandhi, S. M. Miladinović, L.-Y. Cheng, S. Messner, T. Ehrenberger, V. Zanotelli, Y. Butscheid, C. Escher, Mol. Cell. Proteomics 2015, 14, 1400.

[5] S. Tyanova, J. Cox, Cancer systems biology: Methods and protocols 2018, 133.

## References

[1] a) M. Sweigart, C. Zhu, D. Burt, P. DeHoll, C. Agrawal, T. Clanton, K. A. Athanasiou, Ann. Biomed. Eng. 2004, 32, 1569; b) C. A. Murphy, G. M. Cunniffe, A. K. Garg, M. N. Collins, Journal of the mechanical behavior of biomedical materials 2019, 94, 186.

[2] T. K. Tsinman, X. Jiang, L. Han, E. Koyama, R. L. Mauck, N. A. Dyment, FASEB journal: official publication of the Federation of American Societies for Experimental Biology 2021, 35, e21779.

[3] a) A. Abbadessa, J. Crecente-Campo, M. J. Alonso, Tissue Engineering Part B: Reviews 2021, 27, 133; b) J. Twomey-Kozak, C. T. Jayasuriya, Clin. Sports Med. 2020, 39, 125.

[4] F. Porzucek, M. Mankowska, J. A. Semba, P. Cywoniuk, A. Augustyniak, A. M. Mleczko, A. M. Teixeira, P. Martins, A. A. Mieloch, J. D. Rybka, Virtual and Physical Prototyping 2024, 19, e2359620.

[5] J. Pak, J. H. Lee, K. S. Park, J. H. Jeon, S. H. Lee, Open access journal of sports medicine 2017, 33.

[6] M. I. González-Duque, A. M. Flórez, M. A. Torres, M. R. Fontanilla, ACS Biomaterials Science & Engineering 2024, 10, 2426.

[7] J. Schwer, A. Ignatius, A. M. Seitz, Acta Biomaterialia 2023.

[8] H.-W. Yun, B. R. Song, D. I. Shin, X. Y. Yin, M.-D. Truong, S. Noh, Y. J. Jin, H. J. Kwon, B.-H. Min, Materials Science and Engineering: C 2021, 128, 112312.

[9] P. X. Bradley, K. N. Thomas, A. L. Kratzer, A. C. Robinson, J. R. Wittstein, L. E. DeFrate, A. L. McNulty, Curr. Rheumatol. Rep. 2023, 25, 35.

[10] S. Bansal, E. R. Floyd, M. A Kowalski, E. Aikman, P. Elrod, K. Burkey, J. Chahla, R. F. LaPrade, S. A. Maher, J. L. Robinson, Journal of Orthopaedic Research® 2021, 39, 1368.

[11] A. Avila, K. Vasavada, D. S. Shankar, M. Petrera, L. M. Jazrawi, E. J. Strauss, Curr. Rev. Musculoskelet. Med. 2022, 15, 336.

[12] Y. Hashimoto, K. Nishino, K. Orita, S. Yamasaki, Y. Nishida, T. Kinoshita, H. Nakamura, Arthroscopy: The Journal of Arthroscopic & Related Surgery 2022, 38, 441.

[13] J. M. Woodmass, R. F. LaPrade, N. A. Sgaglione, N. Nakamura, A. J. Krych, JBJS 2017, 99, 1222.

[14] M. Keagle, V. Gallicchio, Stem Cells Regen Med 2023, 7, 1.

[15] Z. Li, W. Yan, F. Zhao, H. Wang, J. Cheng, X. Duan, X. Fu, J. Zhang, X. Hu, Y. Ao, Chem. Eng. J. 2023, 473, 145209.

[16] A. Tsujii, N. Nakamura, S. Horibe, The Knee 2017, 24, 1262.

[17] S. Bansal, J. M. Peloquin, N. M. Keah, O. C. O’Reilly, D. M. Elliott, R. L. Mauck, M. H. Zgonis, Journal of Orthopaedic Research® 2020, 38, 2709.

[18] Tissue Engineering Part A 2011, 17, 193.

[19] M. D. Brigham, A. Bick, E. Lo, A. Bendali, J. A. Burdick, A. Khademhosseini, Tissue Engineering Part A 2009, 15, 1645.

[20] M. D. Lombardo, L. Mangiavini, G. M. Peretti, Pharmaceutics 2021, 13, 1886.

[21] S. Park, T. C. Laskow, J. Chen, P. Guha, B. Dawn, D. H. Kim, Aging Cell 2024, e14070.

[22] a) B. Yang, H. Sun, F. Song, M. Yu, Y. Wu, J. Wang, The international journal of biochemistry & cell biology 2017, 87, 104; b) E. Öztürk, E. Despot-Slade, M. Pichler, M. Zenobi-Wong, Exp. Cell Res. 2017, 360, 113; c) S. Dupont, L. Morsut, M. Aragona, E. Enzo, S. Giulitti, M. Cordenonsi, F. Zanconato, J. Le Digabel, M. Forcato, S. Bicciato, Nature 2011, 474, 179; d) W. Zhong, W. Zhang, S. Wang, J. Qin, PLoS One 2013, 8, e61283.

[23] a) A. Karystinou, A. J. Roelofs, A. Neve, F. P. Cantatore, H. Wackerhage, C. De Bari, Arthritis research & therapy 2015, 17, 1; b) W. Zhong, Y. Li, L. Li, W. Zhang, S. Wang, X. Zheng, J. Mol. Histol. 2013, 44, 587; c) W. Zhong, K. Tian, X. Zheng, L. Li, W. Zhang, S. Wang, J. Qin, Stem Cells Dev. 2013, 22, 2083.

[24] H. Alizadeh Sardroud, T. Wanlin, X. Chen, B. F. Eames, Frontiers in Bioengineering and Biotechnology 2022, 9, 787538.

[25] B. Huang, P. Li, M. Chen, L. Peng, X. Luo, G. Tian, H. Wang, L. Wu, Q. Tian, H. Li, Journal of nanobiotechnology 2022, 20, 25.

[26] E. Folkesson, A. Turkiewicz, M. Rydén, H. V. Hughes, N. Ali, J. Tjörnstrand, P. Önnerfjord, M. Englund, Journal of Orthopaedic Research® 2020, 38, 1735.

[27] H.-Y. Ma, Q. Li, W. R. Wong, E.-N. N’Diaye, P. Caplazi, H. Bender, Z. Huang, A. Arlantico, S. Jeet, A. Wong, Science Advances 2023, 9, eadf0133.

[28] W. Kong, C. Lyu, H. Liao, Y. Du, Biomedical Materials 2021, 16, 062005.

[29] B. S. Spearman, N. K. Agrawal, A. Rubiano, C. S. Simmons, S. Mobini, C. E. Schmidt, Journal of Biomedical Materials Research Part A 2020, 108, 279.

[30] H. Kim, B. Kang, X. Cui, S. H. Lee, K. Lee, D. W. Cho, W. Hwang, T. B. Woodfield, K. S. Lim, J. Jang, Advanced Functional Materials 2021, 31, 2011252.

[31] S. Ahn, U. Sharma, K. C. Kasuba, N. Strohmeyer, D. J. Müller, Advanced Science 2023, 10, 2300812.

[32] J. A. Burdick, C. Chung, X. Jia, M. A. Randolph, R. Langer, Biomacromolecules 2005, 6, 386.

[33] S. Chae, S.-S. Lee, Y.-J. Choi, G. Gao, J. H. Wang, D.-W. Cho, Biomaterials 2021, 267, 120466.

[34] R. B. Brackin, G. E. McColgan, S. A. Pucha, M. A. Kowalski, H. Drissi, T. N. Doan, J. M. Patel, Bioengineering 2023, 10, 1013.

[35] a) J. Jang, H.-J. Park, S.-W. Kim, H. Kim, J. Y. Park, S. J. Na, H. J. Kim, M. N. Park, S. H. Choi, S. H. Park, Biomaterials 2017, 112, 264; b) F. Pati, J. Jang, D.-H. Ha, S. Won Kim, J.-W. Rhie, J.-H. Shim, D.-H. Kim, D.-W. Cho, Nature communications 2014, 5, 3935.

[36] D. Szklarczyk, A. Franceschini, S. Wyder, K. Forslund, D. Heller, J. Huerta-Cepas, M. Simonovic, A. Roth, A. Santos, K. P. Tsafou, Nucleic Acids Res. 2015, 43, D447.

[37] P. Shannon, A. Markiel, O. Ozier, N. S. Baliga, J. T. Wang, D. Ramage, N. Amin, B. Schwikowski, T. Ideker, Genome Res. 2003, 13, 2498.

[38] a) D. H. Kim, J. T. Martin, D. M. Elliott, L. J. Smith, R. L. Mauck, Acta Biomaterialia 2015, 12, 21; b) M. Guvendiren, J. A. Burdick, Nature communications 2012, 3, 792.

[39] V. G. Muir, M. Fainor, B. S. Orozco, R. L. Hilliard, M. Boyes, H. E. Smith, R. L. Mauck, T. P. Schaer, J. A. Burdick, S. E. Gullbrand, Advanced Healthcare Materials 2023, 2303326.

[40] K. H. Song, S. J. Heo, A. P. Peredo, M. D. Davidson, R. L. Mauck, J. A. Burdick, Advanced healthcare materials 2020, 9, 1901228.

[41] A. H. Huang, A. Stein, R. S. Tuan, R. L. Mauck, Tissue Engineering Part A 2009, 15, 3461.

[42] a) S.-J. Heo, W. M. Han, S. E. Szczesny, B. D. Cosgrove, D. M. Elliott, D. A. Lee, R. L. Duncan, R. L. Mauck, Biophys. J. 2016, 111, 864; b) R. Mauck, X. Yuan, R. S. Tuan, Osteoarthritis Cartilage 2006, 14, 179.

[43] S. Tarafder, G. Park, C. H. Lee, Connect. Tissue Res. 2020, 61, 292.

