## Supplementary information for "Precision Repair of Zone-Specific Meniscal Injuries Using a Tunable Extracellular Matrix-Based Hydrogel System"

**Supplementary Figures and Legends**

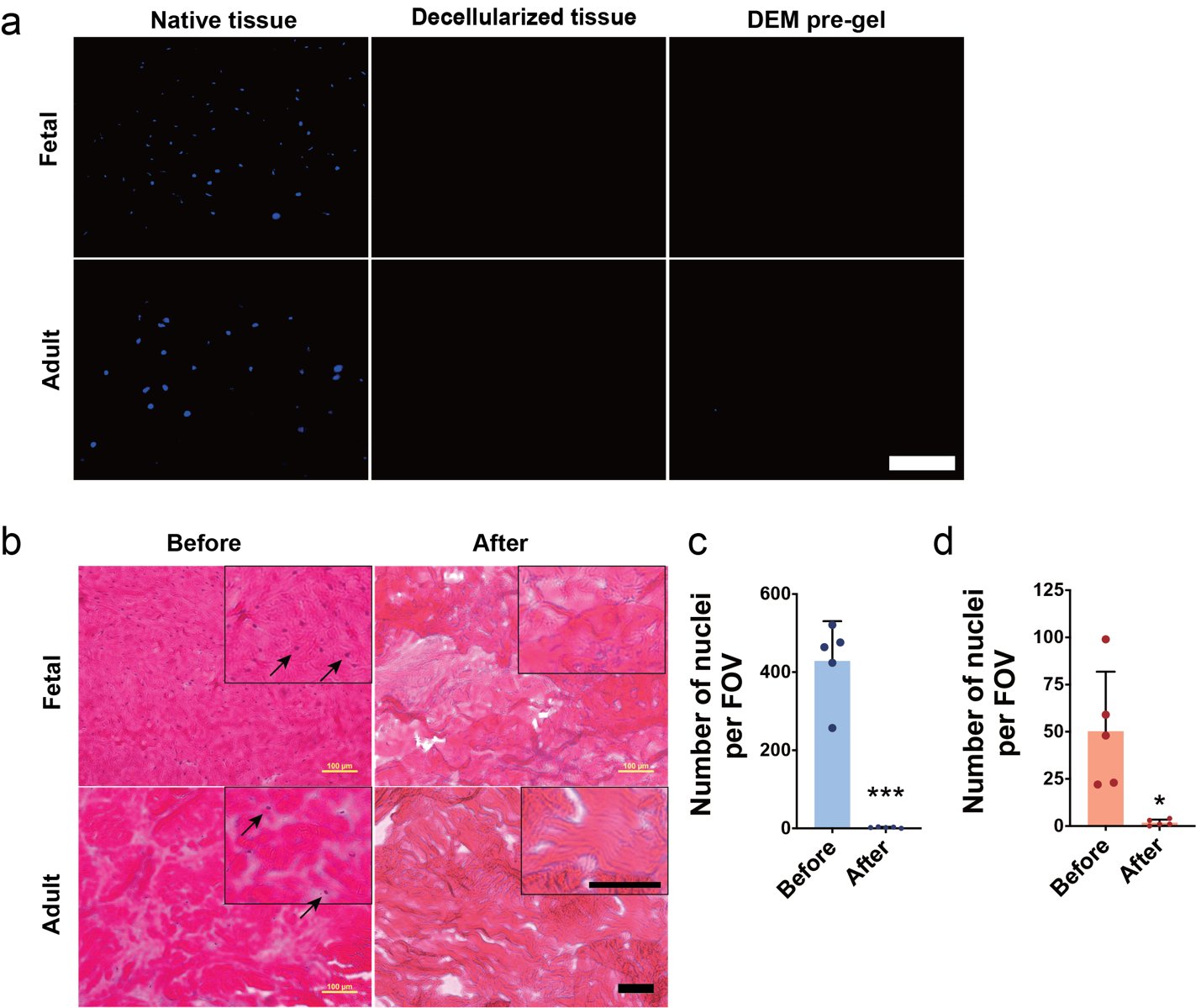

**Supplementary Figure S1**. a) Representative images of DAPI stained tissue sections before and after decellularization and in the pre-gel formulation (scale bar: 100 µm). b) Representative images of H&E stained sections (arrows indicate nuclei; scale bar: 100 µm). Quantification of the number of nuclei per field of view (FOV) in c) Fetal and d) Adult tissue before and after decellularization (n=5; *p<0.05 vs. Before, ***p<0.001 vs. Before).

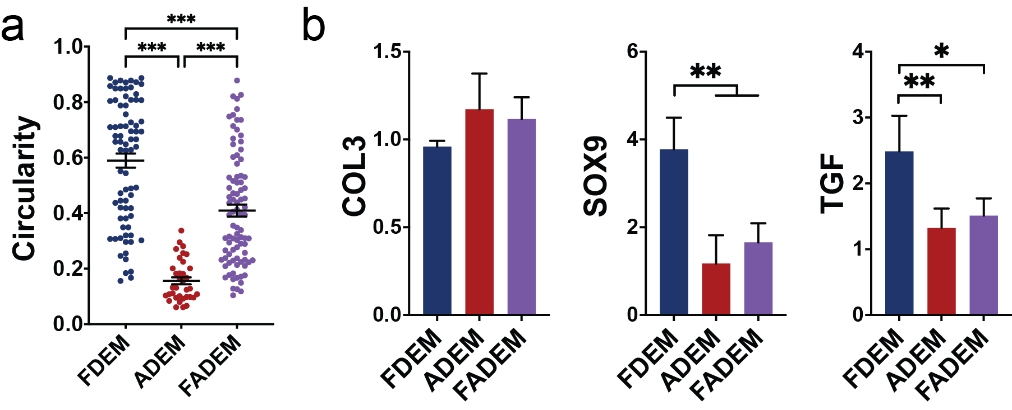

**Supplementary Figure S2.** MSCs in in vitro culture: a) Circularity (n=35-87) on day 3, and b) gene expression on day 7 (n=3-4); *p<0.05, **p<v0.01.

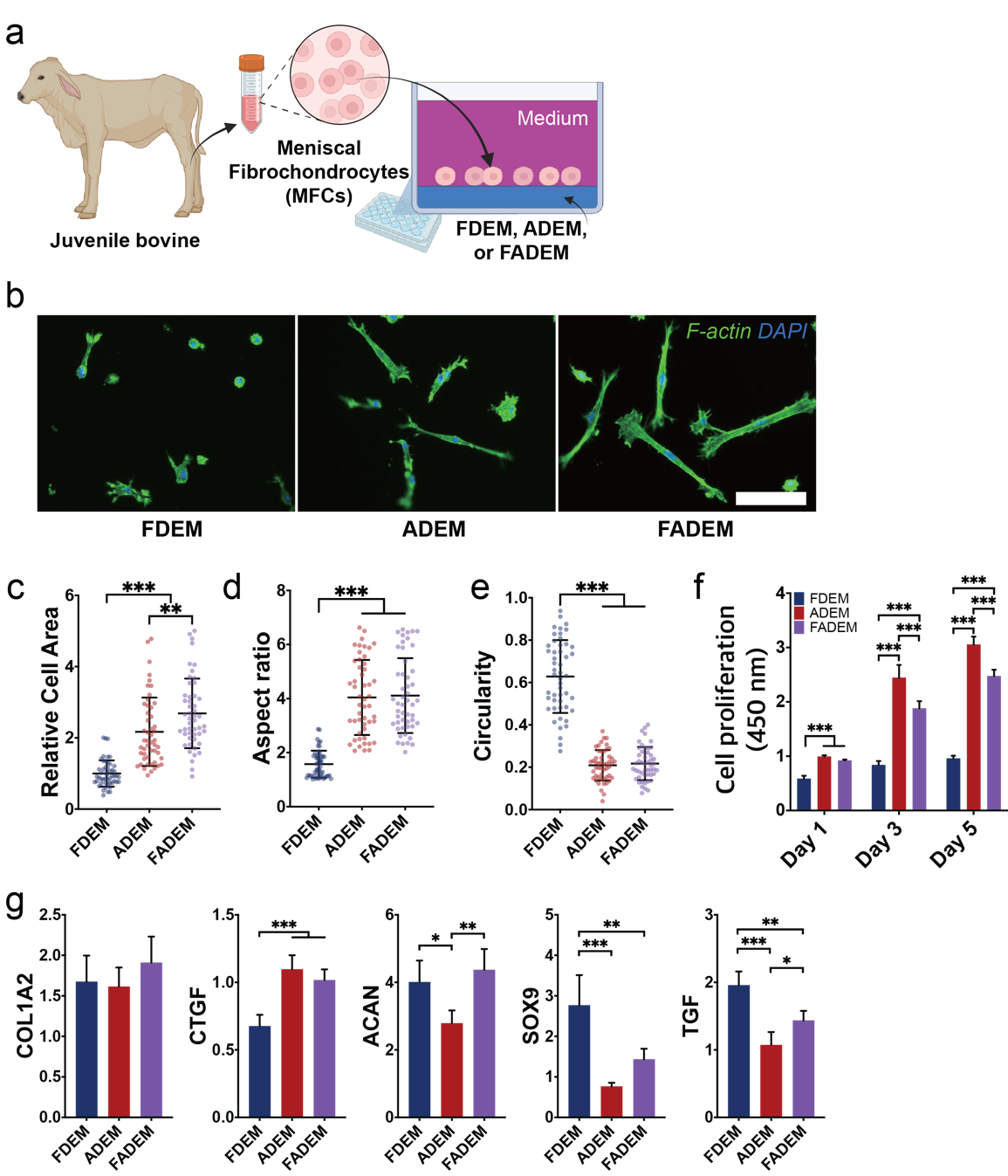

**Supplementary Figure S3.** Meniscus fibrochondrocytes (MFCs) in in vitro culture: a) Illustration of in vitro experiment, b) Representative F-Actin images (Scale bar: 100 µm), c) relative cell area, d) aspect ratio, and e) circularity on Day 3 (n=50); **p<0.01, ***p<0.001). f) Cell proliferation (n=5) and g) gene expression on Day 5 (n=5); COL2 was not detected; **p<0.01, ***p<0.001.

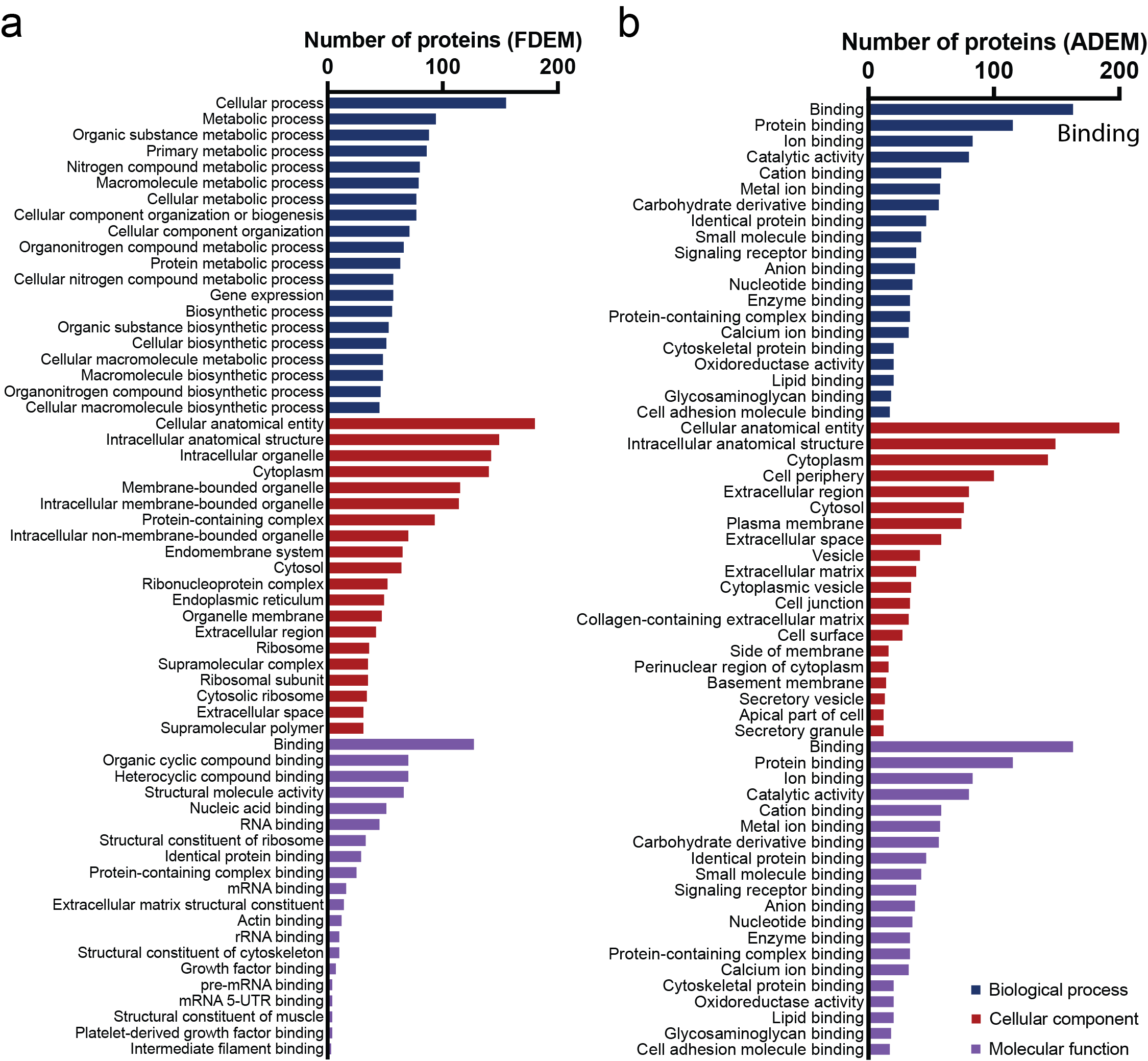

**Supplementary Figure S4.** In-depth gene ontology (GO) analysis for both FDEM and ADEM, derived from four distinct donors. The analysis encompasses: a) FDEM: Showcasing the top 20 GO terms associated with Biological Process, Cellular Component, and Molecular Function, highlighting the cellular activities and interactions prevalent in FDEM, b) ADEM: Similarly, detailing the top 20 GO terms for ADEM, emphasizing the protein, enzyme, and ECM binding activities that are characteristic of this group.

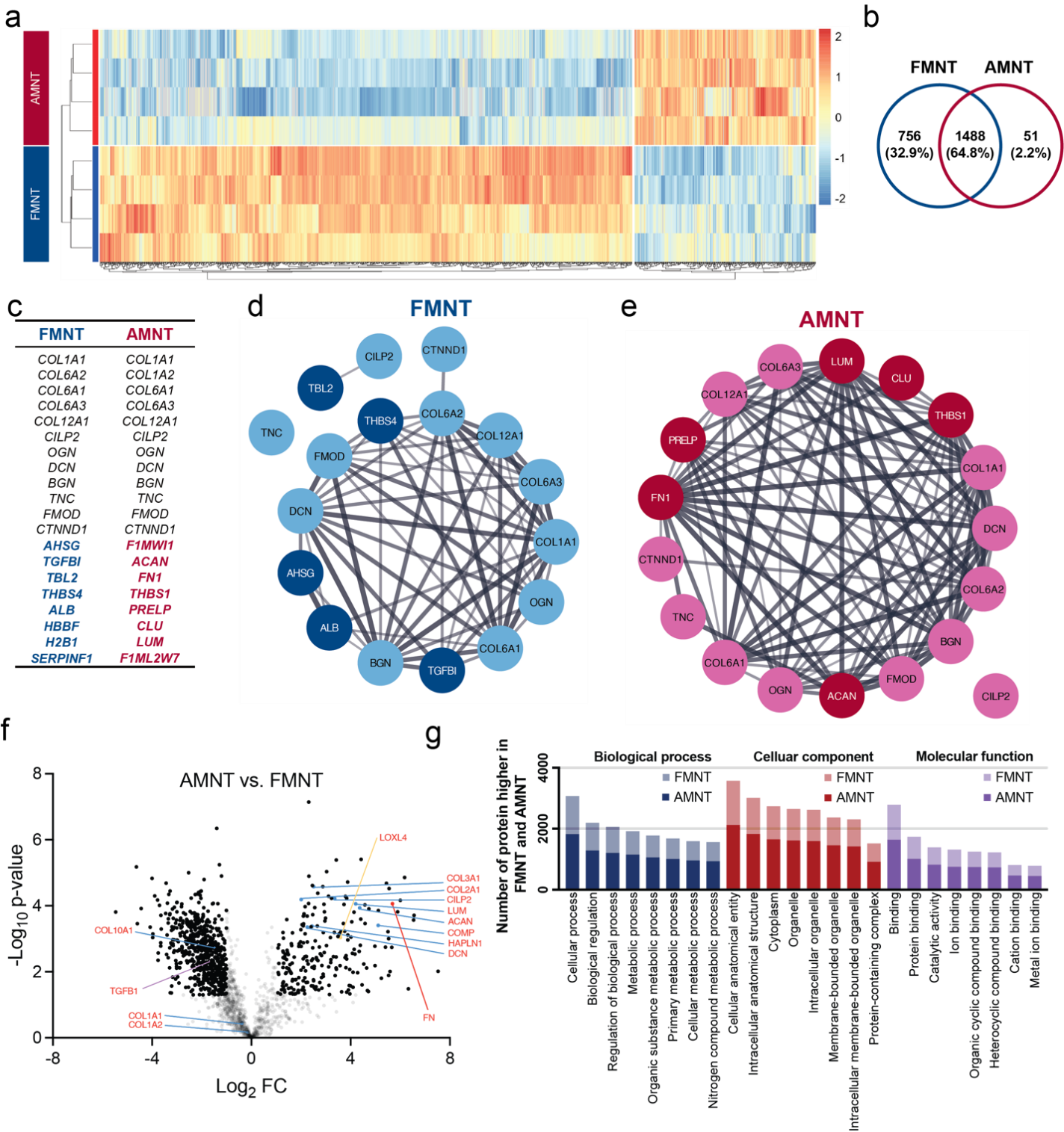

**Supplementary Figure S5**. **Proteomic landscape of native fetal meniscus tissue (FMNT) and native adult meniscus tissue (AMNT), with each group comprising four different donors.** a) Heatmap and clusters demonstrating the protein detection patterns in FMNT and AMNT, indicating the proteomic distinctions between groups, b) Venn diagram illustrating the shared and unique proteins in FMNT and AMNT, c) Listing of top 20 most abundant proteins detected in both native tissues, d-e) Protein-Protein Interaction Networks for the top 20 proteins of FDEM (d) or ADEM (a) with line thickness representing the strength of data support for each interaction, f) Volcano Plot of AMNT (vs. FMNT) showing the differentially abundant proteins between AMNT and FMNT, with statistical significance and fold-change metrics, g) Gene Ontology Analysis indicating the top 8 GO categories for Biological Process, Cellular Component, and Molecular Function, with color-coding to differentiate the categories that vary between FMNT and AMNT.

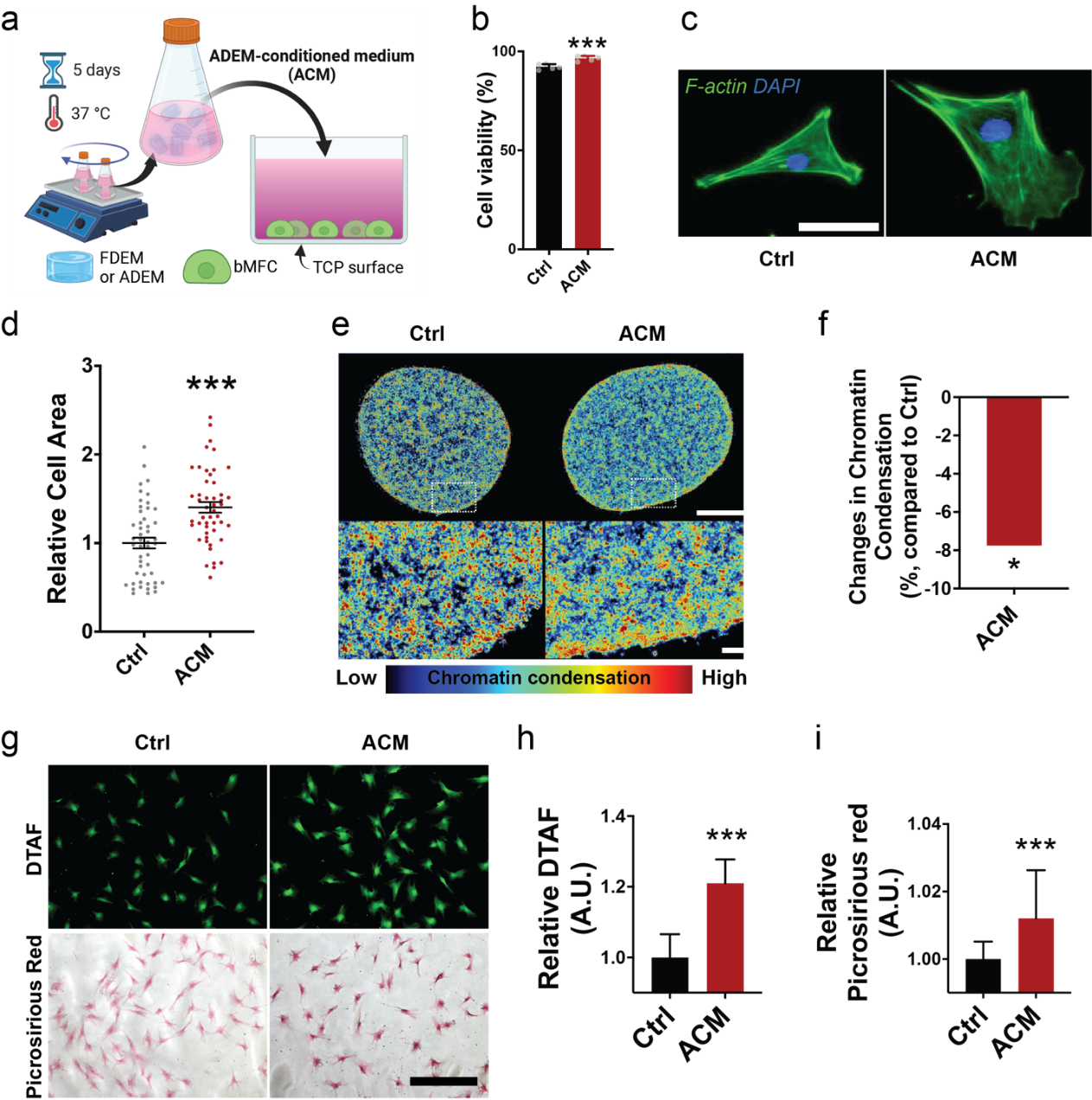

**Supplementary Figure S6**. a) Schematic of experiment setup for the conditioned media assay, b) Cytotoxicity results after 5 days (n=5; ***: p<0.001), c) Representative F-actin images from day 3 (Scale bar: 50 μm), d) Quantified cell area (n=46-49; ***: p<0.001 vs. Ctrl), (e) Representative images showing chromatin condensation levels in MFCs [Scale bars: 1 μm (top) and 500 nm (bottom)], and f) Chromatin condensation quantification (*: p<0.05 vs. Ctrl). Protein staining: g) Representative images of DTAF staining and Picrosirius Red staining (Scale bar: 200 um) of MFCs. Relative intensity measurements of the DTAF (h) and PSR (i) staining intensity per cell (n=25; *: p<0.05 vs. Ctrl, ***: p<0.001 vs. Ctrl).

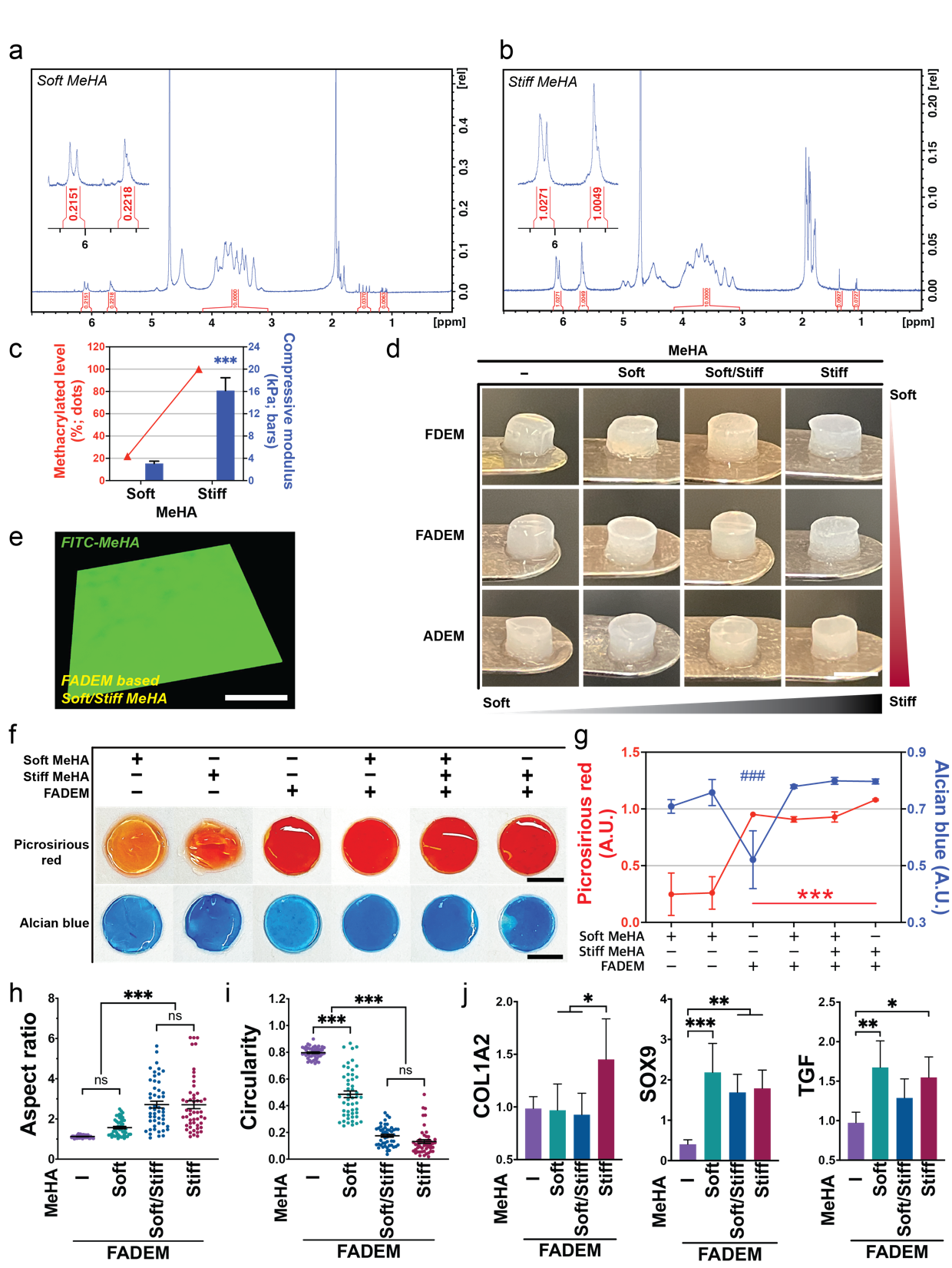

**Supplementary Figure S7.** a) Degree of methacrylation analysis by ¹H-NMR: Soft MeHA. b) Stiff MeHA.

c) % Methacrylation and Mechanical Properties: Comparison of % methacrylation and mechanical properties of MeHA hydrogels (n=5; ***p<0.001 vs. Stiff MeHA), d) Images of fabricated stiffness tunable HA/DEM hydrogels (scale bars: 3 mm). e) Fluorescence images showing homogeneity: ‘soft/stiff’ FADEM-based hydrogels (Green: ‘soft/stiff MeHA distribution; Scale bar: 400 µm), f) Collagen and proteoglycan staining: representative images of Picrosirius red (PSR) and Alcian blue (AB) staining, g) Quantification of PSR and AB staining intensity (n=5; ###: p<0.001 vs. whole groups in Alcian blue graph; ***: p<0.001 vs. Soft MeHA in PSR graph). Quantification of cell aspect ratio (h) and circularity (i) (n=50; ***p<0.001). j) Quantification of gene expression by MSCs on stiffness-tunable HA/DEM hydrogels (n=4-5; *p<0.05, **p<0.01, ***p<0.001).

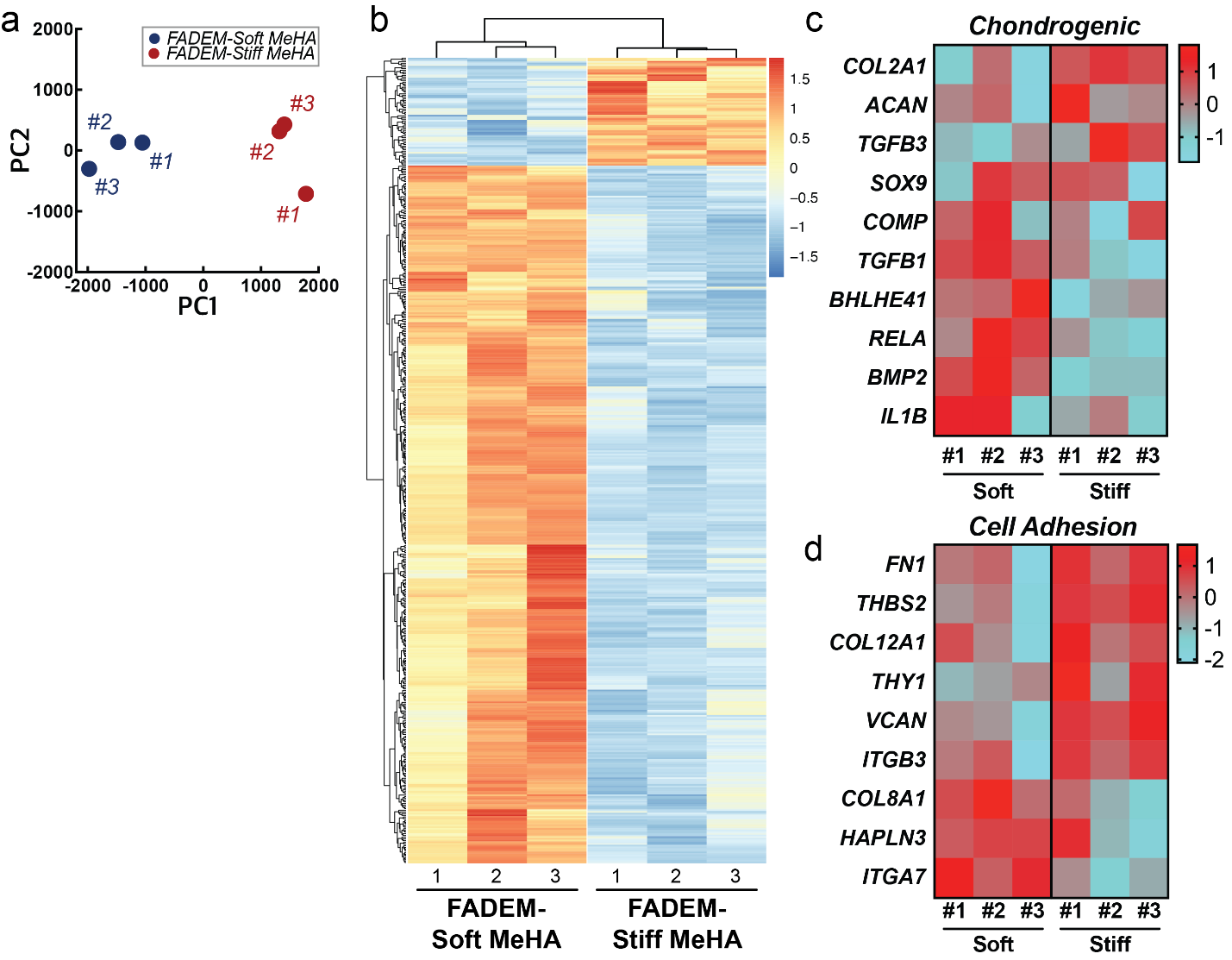

**Supplementary Figure S8.** a) PCA plot clustering of samples, b) Heatmap showing expression patterns of significantly altered genes across different experimental conditions. Z-score heat maps of Chondrogenic (c) and cell adhesion (d) related gene expression (n=3/group).

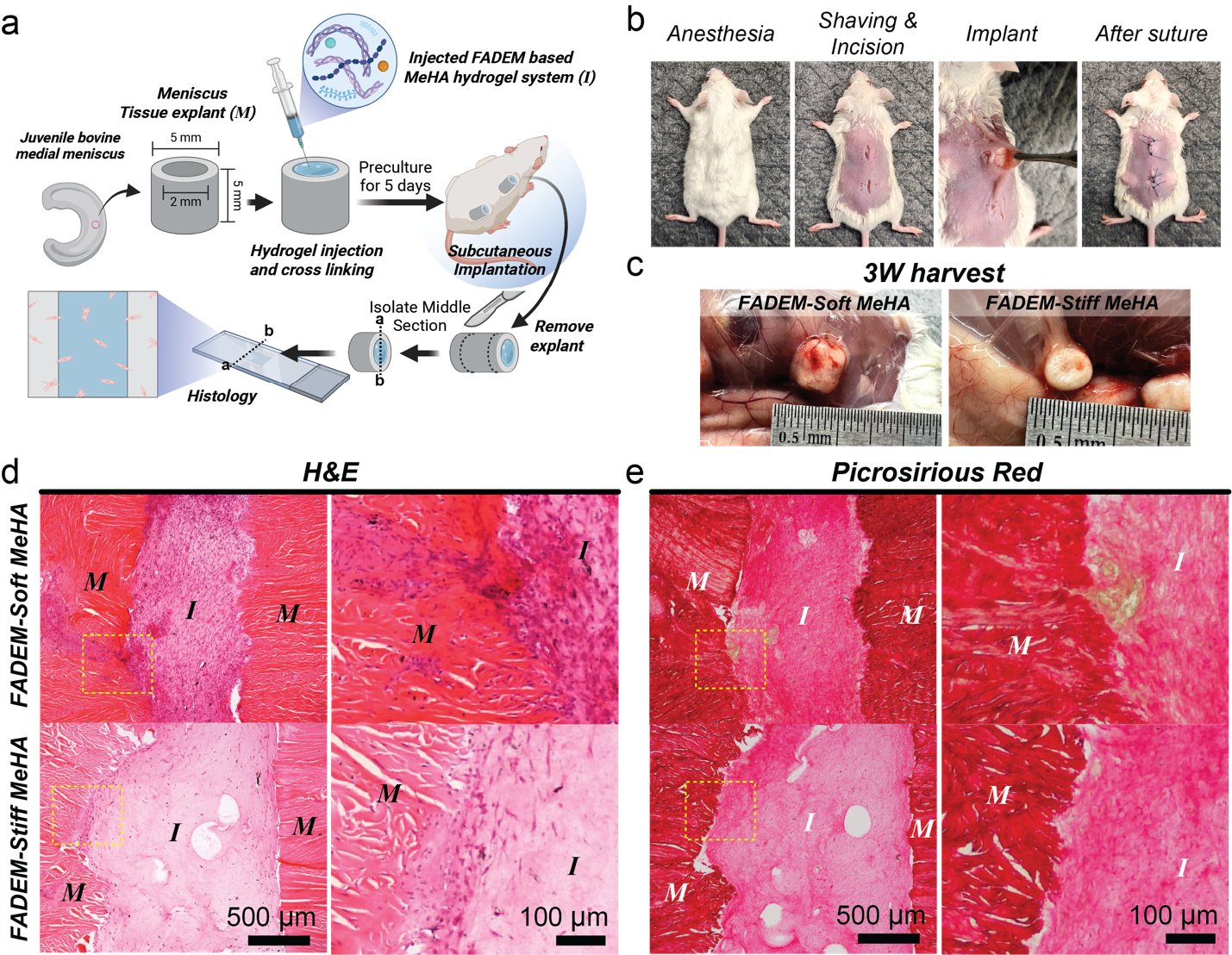

**Supplementary Figure S9**. a) Schematic illustration of bovine tissue implant with injected FADEM-based tunable MeHA hydrogel, subcutaneous implantation in mice, and preparation for histological staining, b) Images showcasing the surgical procedure, c) Harvested samples at 3 weeks post-implantation. Histological Analysis (3 weeks post-implantation): d) Representative H&E-stained and e) Picrosirius red-stained sections (M: Meniscus tissue explant, I: Injected FADEM-based MeHA hydrogel system).

**Supplementary Table 1. Top 30 up- or down-regulated genes**

| **FADEM-Stiff MeHA vs. FADEM-Soft MeHA** | | | | | |
| --- | --- | --- | --- | --- | --- |
| **Downregulated** | | | **Upregulated** | | |
| **Symbol** | **Name** | **Fold**  **(Log2)** | **Symbol** | **Name** | **Fold**  **(Log2)** |
| MATK | Megakaryocyte-associated tyrosine kinase | -6.32 | ZNF711 | Zinc finger protein 711 | 3.07 |
| MRPL36 | Mitochondrial ribosomal protein L36 | -4.34 | CRISPLD1 | Cysteine rich secretory protein LCCL domain containing 1 | 2.57 |
| CITED4 | Cbp/p300 interacting transactivator with Glu/Asp rich carboxy-terminal domain 4 | -4.30 | DKK2 | Dickkopf WNT signaling pathway inhibitor 2 | 2.54 |
| C6H4orf48 | Chromosome 6 C4orf48 homolog | -4.30 | SST | Somatostatin | 2.53 |
| KCTD12 | Potassium channel tetramerization domain containing 12 | -4.06 | SVEP1 | Sushi, von Willebrand factor type A, EGF and pentraxin domain containing 1 | 2.53 |
| MAFB | MAF bZIP transcription factor B | -4.05 | IGDCC4 | Immunoglobulin superfamily DCC subclass member 4 | 2.45 |
| NACC2 | NACC family member 2 | -3.88 | ADAMTS1 | ADAM metallopeptidase with thrombospondin type 1 motif 1 | 2.36 |
| INAFM1 | InaF motif containing 1 | -3.87 | INHBB | Inhibin subunit beta B | 2.15 |
| PCSK1N | Proprotein convertase subtilisin/kexin type 1 inhibitor | -3.87 | LIPG | lipase G, endothelial type | 2.03 |
| MYLK2 | Myosin light chain kinase 2 | -3.54 | FAM13C | Family with sequence similarity 13 member C | 2.00 |
| COL11A2 | Collagen type XI alpha 2 chain | -3.41 | IGFBP5 | Insulin like growth factor binding protein 5 | 1.97 |
| TUSC1 | Tumor suppressor candidate 1 | -3.40 | MALAT1 | Metastasis associated lung adenocarcinoma transcript 1 | 1.90 |
| IFITM5 | Interferon induced transmembrane protein 5 | -3.35 | CNTNAP4 | Contactin associated protein family member 4 | 1.86 |
| TMEM203 | Transmembrane protein 203 | -3.18 | MMP13 | Matrix metallopeptidase 13 | 1.74 |
| PNPLA2 | Patatin like phospholipase domain containing 2 | -3.15 | FAT4 | FAT atypical cadherin 4 | 1.68 |
| FAM229A | Family with sequence similarity 229 member A | -3.13 | DDAH1 | Dimethylarginine dimethylaminohydrolase 1 | 1.67 |
| MRM1 | Mitochondrial rRNA methyltransferase 1 | -3.11 | ATP13A3 | ATPase 13A3 | 1.64 |
| SFRP5 | Secreted frizzled related protein 5 | -3.05 | COL2A1 | Collagen type II alpha 1 chain | 1.64 |
| METRN | Meteorin, glial cell differentiation regulator | -2.95 | KIAA0408 | KIAA0408 | 1.61 |
| CD81 | CD81 molecule | -2.95 | VGLL3 | Vestigial like family member 3 | 1.60 |
| FNDC4 | Fibronectin type III domain containing 4 | -2.90 | PPM1K | Protein phosphatase, Mg2+/Mn2+ dependent 1K | 1.52 |
| LGI2 | Leucine rich repeat LGI family member 2 | -2.90 | MATN2 | Matrilin 2 | 1.50 |
| NDRG4 | NDRG family member 4 | -2.89 | EPHA4 | EPH receptor A4 | 1.45 |
| SNAP29 | Synaptosome associated protein 29 | -2.88 | RBL1 | RB transcriptional corepressor like 1 | 1.42 |
| NAT14 | N-acetyltransferase 14 (putative) | -2.83 | EFEMP1 | EGF containing fibulin extracellular matrix protein 1 | 1.39 |
| FZD5 | Frizzled class receptor 5 | -2.79 | NID1 | Nidogen 1 | 1.39 |
| MOB2 | MOB kinase activator 2 | -2.79 | ADAMTSL3 | ADAMTS like 3 | 1.39 |
| ZXDC | ZXD family zinc finger C | -2.77 | CCNJ | Cyclin J | 1.35 |
| MLYCD | Malonyl-CoA decarboxylase | -2.77 | MAN1A1 | Mannosidase alpha class 1A member 1 | 1.34 |
| RAPGEF3 | Rap guanine nucleotide exchange factor | -2.77 | XIRP1 | Xin actin binding repeat containing 1 | 1.31 |

**Supplementary Table 2. Tukey's multiple comparisons test for stiffness of mechanical modulated DEM-based MeHA hydrogel system (Figure 5b; n=5; ns: not significant, *<0.05, **p<0.01, ***p<0.001)**

| **Tukey's multiple comparisons test (H: MeHA)** | | | | | |
| --- | --- | --- | --- | --- | --- |
| **Comparisons** | **Adjusted P Value** | **Summary** | **Comparisons** | **Adjusted P Value** | **Summary** |
| FDEM vs. FDEM-Soft H | 0.9817 | ns | FDEM-Stiff H vs. Stiff H | 0.0953 | ns |
| FDEM vs. FDEM-Soft/Stiff H | <0.0001 | *** | FADEM vs. FADEM-Soft H | 0.0844 | ns |
| FDEM vs. FDEM-Stiff H | <0.0001 | *** | FADEM vs. FADEM-Soft/Stiff H | <0.0001 | *** |
| FDEM vs. FADEM | 0.9971 | ns | FADEM vs. FADEM-Stiff H | <0.0001 | *** |
| FDEM vs. FADEM-Soft H | 0.0033 | ** | FADEM vs. ADEM | 0.0354 | * |
| FDEM vs. FADEM-Soft/Stiff H | <0.0001 | *** | FADEM vs. ADEM-Soft H | <0.0001 | *** |
| FDEM vs. FADEM-Stiff H | <0.0001 | *** | FADEM vs. ADEM-Soft/Stiff H | <0.0001 | *** |
| FDEM vs. ADEM | 0.0011 | ** | FADEM vs. ADEM-Stiff H | <0.0001 | *** |
| FDEM vs. ADEM-Soft H | <0.0001 | *** | FADEM vs. Soft H | 0.3775 | ns |
| FDEM vs. ADEM-Soft/Stiff H | <0.0001 | *** | FADEM vs. Stiff H | <0.0001 | *** |
| FDEM vs. ADEM-Stiff H | <0.0001 | *** | FADEM-Soft H vs. FADEM-Soft/Stiff H | <0.0001 | *** |
| FDEM vs. ADEM-Soft/Stiff H | 0.9691 | ns | FADEM-Soft H vs. FADEM-Stiff H | <0.0001 | *** |
| FDEM vs. ADEM-Stiff H | <0.0001 | *** | FADEM-Soft H vs. ADEM | >0.9999 | ns |
| FDEM-Soft H vs. FDEM-Soft/Stiff H | 0.0007 | *** | FADEM-Soft H vs. ADEM-Soft H | 0.2375 | ns |
| FDEM-Soft H vs. FDEM-Stiff H | <0.0001 | *** | FADEM-Soft H vs. ADEM-Soft/Stiff H | <0.0001 | *** |
| FDEM-Soft H vs. FADEM | >0.9999 | ns | FADEM-Soft H vs. ADEM-Stiff H | <0.0001 | *** |
| FDEM-Soft H vs. FADEM-Soft H | 0.1526 | ns | FADEM-Soft H vs. Soft H | <0.0001 | *** |
| FDEM-Soft H vs. FADEM-Soft/Stiff H | <0.0001 | *** | FADEM-Soft H vs. Stiff H | <0.0001 | *** |
| FDEM-Soft H vs. FADEM-Stiff H | <0.0001 | *** | FADEM-Soft/Stiff H vs. FADEM-Stiff H | <0.0001 | *** |
| FDEM-Soft H vs. ADEM | 0.0692 | ns | FADEM-Soft/Stiff H vs. ADEM | <0.0001 | *** |
| FDEM-Soft H vs. ADEM-Soft H | <0.0001 | *** | FADEM-Soft/Stiff H vs. ADEM-Soft H | 0.0908 | ns |
| FDEM-Soft H vs. ADEM-Soft/Stiff H | <0.0001 | *** | FADEM-Soft/Stiff H vs. ADEM-Soft/Stiff H | 0.0005 | *** |
| FDEM-Soft H vs. ADEM-Stiff H | <0.0001 | *** | FADEM-Soft/Stiff H vs. ADEM-Stiff H | <0.0001 | *** |
| FDEM-Soft H vs. Soft H | 0.2397 | ns | FADEM-Soft/Stiff H vs. Soft H | <0.0001 | *** |
| FDEM-Soft H vs. Stiff H | <0.0001 | *** | FADEM-Soft/Stiff H vs. Stiff H | 0.4774 | ns |
| FDEM-Soft/Stiff H vs. FDEM-Stiff H | <0.0001 | *** | FADEM-Stiff H vs. ADEM | <0.0001 | *** |
| FDEM-Soft/Stiff H vs. FADEM | 0.0003 | *** | FADEM-Stiff H vs. ADEM-Soft H | <0.0001 | *** |
| FDEM-Soft/Stiff H vs. FADEM-Soft H | 0.8557 | ns | FADEM-Stiff H vs. ADEM-Soft/Stiff H | <0.0001 | *** |
| FDEM-Soft/Stiff H vs. FADEM-Soft/Stiff H | 0.0051 | ** | FADEM-Stiff H vs. ADEM-Stiff H | 0.9661 | ns |
| FDEM-Soft/Stiff H vs. FADEM-Stiff H | <0.0001 | *** | FADEM-Stiff H vs. Soft H | <0.0001 | *** |
| FDEM-Soft/Stiff H vs. ADEM | 0.9636 | ns | FADEM-Stiff H vs. Stiff H | <0.0001 | *** |
| FDEM-Soft/Stiff H vs. ADEM-Soft H | 0.9989 | ns | ADEM vs. ADEM-Soft H | 0.4246 | ns |
| FDEM-Soft/Stiff H vs. ADEM-Soft/Stiff H | <0.0001 | *** | ADEM vs. ADEM-Soft/Stiff H | <0.0001 | *** |
| FDEM-Soft/Stiff H vs. ADEM-Stiff H | <0.0001 | *** | ADEM vs. ADEM-Stiff H | <0.0001 | *** |
| FDEM-Soft/Stiff H vs. Soft H | <0.0001 | *** | ADEM vs. Soft H | <0.0001 | *** |
| FDEM-Soft/Stiff H vs. Stiff H | <0.0001 | *** | ADEM vs. Stiff H | <0.0001 | *** |
| FDEM-Stiff H vs. FADEM | <0.0001 | *** | ADEM-Soft H vs. ADEM-Soft/Stiff H | <0.0001 | *** |
| FDEM-Stiff H vs. FADEM-Soft H | <0.0001 | *** | ADEM-Soft H vs. ADEM-Stiff H | <0.0001 | *** |
| FDEM-Stiff H vs. FADEM-Soft/Stiff H | <0.0001 | *** | ADEM-Soft H vs. Soft H | <0.0001 | *** |
| FDEM-Stiff H vs. FADEM-Stiff H | <0.0001 | *** | ADEM-Soft H vs. Stiff H | <0.0001 | *** |
| FDEM-Stiff H vs. ADEM | <0.0001 | *** | ADEM-Soft/Stiff H vs. ADEM-Stiff H | <0.0001 | *** |
| FDEM-Stiff H vs. ADEM-Soft H | <0.0001 | *** | ADEM-Soft/Stiff H vs. Soft H | <0.0001 | *** |
| FDEM-Stiff H vs. ADEM-Soft/Stiff H | >0.9999 | ns | ADEM-Soft/Stiff H vs. Stiff H | 0.3756 | ns |
| FDEM vs. FDEM-Soft H | 0.9817 | ns | ADEM-Stiff H vs. Soft H | <0.0001 | *** |
| FDEM-Stiff H vs. ADEM-Stiff H | <0.0001 | *** | ADEM-Stiff vs. Stiff H | <0.0001 | *** |
| FDEM-Stiff H vs. Soft H | <0.0001 | *** | Soft H vs. Stiff H | <0.0001 | *** |

### **Supplementary References**

[1]	A. Naba, K. R. Clauser, R. O. Hynes, *JoVE (Journal of Visualized Experiments)* **2015**, e53057.

[2]	S. Tyanova, T. Temu, J. Cox, *Nat. Protoc.* **2016**, 11, 2301.

[3]	C. Bielow, G. Mastrobuoni, S. Kempa, *J. Proteome Res.* **2016**, 15, 777.

[4]	R. Bruderer, O. M. Bernhardt, T. Gandhi, S. M. Miladinović, L.-Y. Cheng, S. Messner, T. Ehrenberger, V. Zanotelli, Y. Butscheid, C. Escher, *Mol. Cell. Proteomics* **2015**, 14, 1400.

[5]	S. Tyanova, J. Cox, *Cancer systems biology: Methods and protocols* **2018**, 133.
